# Conserved human effector Treg signature is reflected in transcriptomic and epigenetic landscape

**DOI:** 10.1101/2020.09.30.319962

**Authors:** Gerdien Mijnheer, Lisanne Lutter, Michal Mokry, Marlot van der Wal, Veerle Fleskens, Rianne Scholman, Aridaman Pandit, Weiyang Tao, Mark Wekking, Stephin Vervoort, Ceri Roberts, Alessandra Petrelli, Janneke G.C. Peeters, Marthe Knijff, Sytze de Roock, Sebastiaan Vastert, Leonie S. Taams, Jorg van Loosdregt, Femke van Wijk

**Author notes:** Shared position (Gerdien Mijnheer, Lisanne Lutter, Jorg van Loosdregt, Femke van Wijk). Corresponding author: Femke van Wijk, PhD, Center for Translational Immunology, Wilhelmina Children’s Hospital, University Medical Center Utrecht, Lundlaan 6, 3584 EA Utrecht.

## Abstract

Treg are critical regulators of immune homeostasis, and increasing evidence demonstrates that environment-driven Treg differentiation into effector (e)Treg is crucial for optimal functioning. However, human Treg programming under inflammatory conditions remains poorly understood. Here, we combine transcriptional and epigenetic profiling to identify the human eTreg core signature. Functional autoimmune inflammation-derived Treg display a unique transcriptional profile characterized by upregulation of both a core Treg (FOXP3, CTLA-4, TIGIT) and effector program (GITR, BLIMP-1, BATF). We identified a specific human eTreg signature that includes the vitamin D receptor (VDR) as predicted key-regulator in eTreg differentiation. H3K27ac/H3K4me1 occupancy revealed pronounced changes in the (super-)enhancer landscape, including enrichment of the binding motif for VDR and BATF. The observed Treg profile showed striking overlap with tumor-infiltrating Treg. Our data demonstrate that human inflammation-derived Treg acquire a specific eTreg profile guided by epigenetic changes. The core eTreg profile is conserved, and fine-tuned by environment-specific adaptations.

## Introduction

Forkhead box P3-expressing (FOXP3^+^) Treg are key players to control aberrant immune responses. Mutations in the *FOXP3* gene lead to severe autoimmunity and inflammation in both mice and men^1,2^. Because of their potential for clinical applications, Treg have been intensively studied over the last decades. Despite this effort there is still a fundamental gap in knowledge, especially regarding the gene expression profiles and epigenetic regulation of human Treg in inflammatory settings.

There is accumulating evidence from mouse models that specific environments such as inflammation or non-lymphoid tissues can induce further differentiation/adaptation into specialized activated Treg subsets, also referred to as effector (e)Treg^3,4^ (reviewed in ref. ^5^). Characteristic for eTreg appears to be the maintenance of FOXP3 expression, increased expression of several molecules related to their function such as ICOS, CTLA4 and TIGIT, and adaptation to the local environment. For example, recent studies in mice show that in visceral adipose tissue Treg play a dominant role in metabolic homeostasis^6^, whereas Treg in skin and gut can promote wound repair and are involved in local stem cell maintenance^7,8^. It remains poorly understood how this specialization in tissues is acquired and regulated at a transcriptional and epigenetic level.

In mice, eTreg have been demonstrated to be crucial in specific inflammatory settings. Interestingly, inflammatory signals can induce the stable upregulation of typical Th cell transcription factors such as T-bet, allowing the Treg to migrate to the site of inflammation^9^. Moreover, co-expression of T-bet and Foxp3 is essential to prevent severe Th1 autoimmunity^10^. Studies using transgenic mice have eminently contributed to the knowledge in this field, but how this can be translated to humans remains to be elucidated. Because of the interest in the use of Treg for therapeutic purposes in a variety of human diseases it is highly relevant to gain insight in human eTreg programming in inflammatory settings.

Also in humans there is evidence of environment-induced adaptation of human (e)Treg^4,7,11^ (reviewed in ref. ^12^) including tumor-infiltrating Treg signatures^13–15^. Pertinent issues that remain to be addressed are how diverse and stable human eTreg programming is in the inflammatory environment, and whether this is regulated at an epigenetic level.

Here, we investigated the gene expression profile and active enhancer landscape of human Treg in an autoimmune-associated inflammatory environment, i.e. synovial fluid (SF) from inflamed joints of Juvenile Idiopathic Arthritis (JIA) and Rheumatoid Arthritis (RA) patients. Transcriptome analysis of SF Treg revealed a classical Treg signature as well as strongly upregulated markers including ICOS, BATF, T-bet, IL-12rβ2 and Blimp-1, indicating eTreg differentiation and adaptation to the inflammatory environment. Data-driven network analysis further revealed the vitamin D_3_ receptor (VDR) as a predicted key-regulator of eTreg differentiation. Importantly, H3K27ac and H3K4me1 ChIP-seq showed that this dual programming was imprinted at an epigenetic level. These eTreg demonstrated normal suppressive function and enhanced IL-2 signaling. Finally, a substantial overlap between the SF Treg signature and recently published human tumor-infiltrating Treg signatures were found. These findings indicate that human eTreg programming is epigenetically imprinted and may be commonly induced in inflammatory conditions ranging from autoimmune to tumor settings.

## Results

### Suppressive Treg from human inflamed joint fluid demonstrate a distinct transcriptional profile with an enhanced core Treg signature

To determine whether the gene expression profile of human Treg in an inflammatory environment is different from circulating human Treg, CD3^+^CD4^+^CD25^+^CD127^low^ Treg and CD3^+^CD4^+^CD25^−^CD127^+^ non-Treg were isolated from: synovial exudate obtained from inflamed joints of JIA patients, peripheral blood (PB) from JIA patients with active and inactive disease, and PB from healthy children and healthy adults (Supplementary Fig. 1a for gating strategy). More than 90% of the sorted Treg populations were FOXP3 positive (Supplementary Fig. 1b). The transcriptional landscape was determined with RNA-sequencing. An unsupervised principal component analysis (PCA) was performed to study the variability between Treg derived from different environments. SF-derived Treg clearly clustered separately from PB-derived Treg (Fig. 1a), indicating that SF-derived Treg have a specific expression pattern compared to PB-derived Treg.

**Fig. 1.**
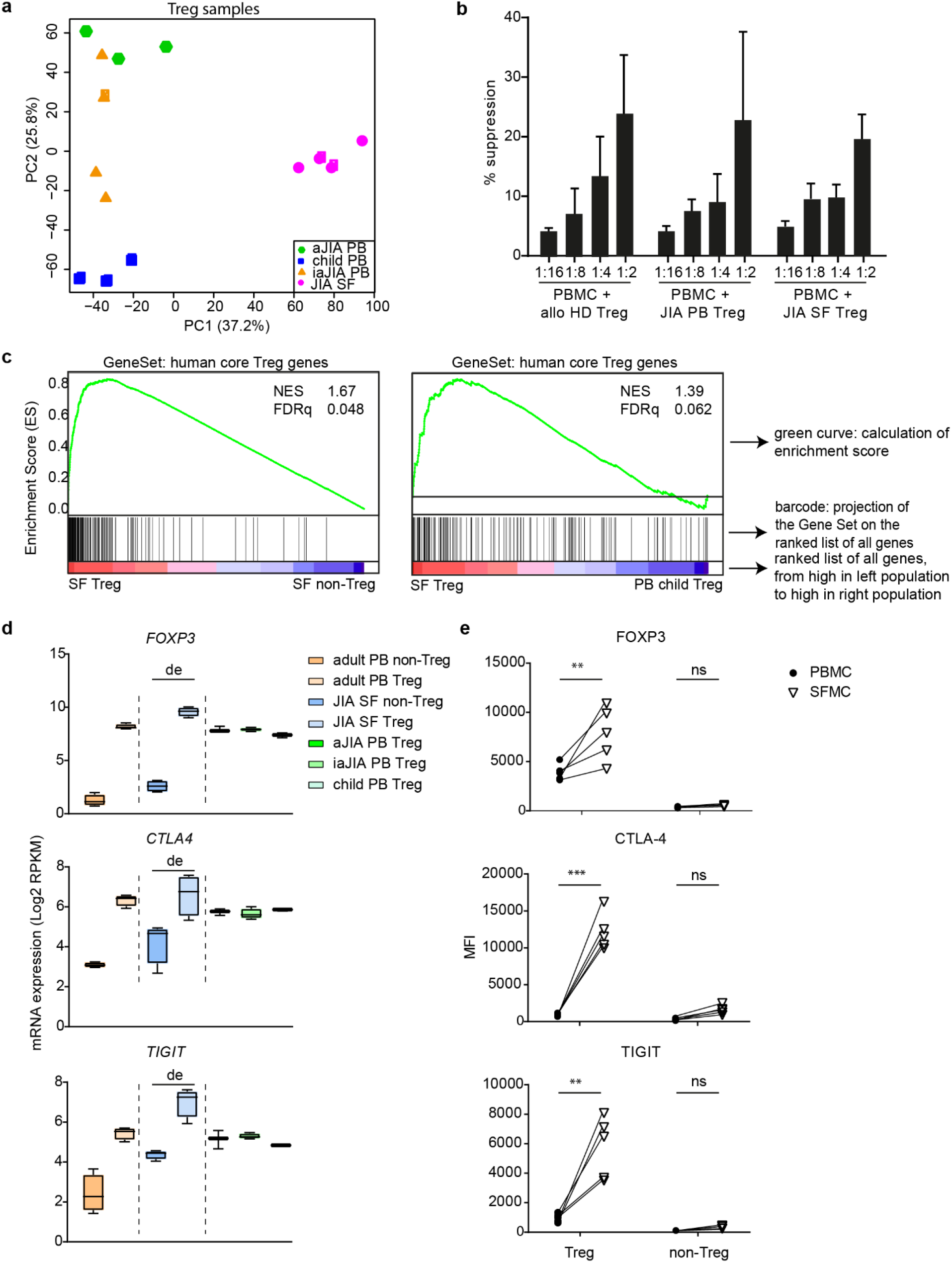
Human inflammation-derived Treg are distinct from peripheral Treg with clear Treg characteristics. **a.** Unsupervised Principal Component Analysis (PCA) of all Treg groups, synovial fluid (SF)- and peripheral blood (PB)-derived, from children. **b** Bar charts representing suppression (percentage) of CD4^+^ T cells by Treg from PB of healthy donors or juvenile idiopathic arthritis (JIA) patients and SF of JIA patients. 50.000 ctViolet labeled PBMC from an allogenic healthy donor were cultured with different ratios of Treg for 4 days in anti-CD3 coated plates (n = 4, mean + SEM). **c** GeneSet Enrichment Analysis (GSEA) of human core Treg signature genes (identified by Ferraro *et al.*^65^) in pairwise comparisons involving SF Treg and non-Treg, and SF Treg and healthy child PB Treg, represented by the normalized enrichment score (NES) and FDR statistical value (FDRq). **d** mRNA expression (log2 RPKM) of *FOXP3*, *CTLA4* and *TIGIT* in Treg derived from PB of healthy adults (adult), healthy children (child), JIA patients with active (aJIA) or inactive (iaJIA) disease, SF of JIA patients (JIA SF) and non-Treg from PB of healthy adults and SF of JIA patients (de = differentially expressed according to log2 fold change ≥ 0.6, adjusted (adj)p-value ≤ 0.05, mean of all normalized counts > 10; adj p-values *FOXP3* = 1.1E^−95^, *CTLA4* = 2.3E^−05^, *TIGIT* = 1.9E^−17^) **e** Median Fluorescence Intensity (MFI) of FOXP3, CTLA-4 and TIGIT in CD3^+^CD4^+^CD25^+^CD127^low^ Treg and CD3^+^CD4^+^CD25^−^CD127^+^ non-Treg from paired SFMC and PBMC from 5 JIA patients. Statistical comparisons were performed using two-way ANOVA with Sidak correction for multiple testing: ns, \*\**P* < 0.01, \*\*\**P* < 0.001. (**b** and **e**) Data are representative of two independent experiments.

In accordance with previous publications we confirmed that SF-derived Treg were functional using an *in vitro* suppression assay (Fig. 1b)^16,17^. As expected, Treg signature genes were significantly enriched in SF Treg compared to SF CD4^+^ non-Treg (Fig. 1c, left panel). Importantly, Treg signature genes were also enriched in SF compared to PB Treg derived from both healthy children and JIA patients (Fig. 1c right panel and Supplementary Fig. 2a). These included Treg hallmark genes important for Treg stability and function such as *FOXP3*, *CTLA4* and *TIGIT*, at both the transcriptional and protein level (Fig. 1d, e and Supplementary Fig. 1a and 2b)^18,19^. In humans, CD3^+^CD4^+^non-Treg can upregulate FOXP3 and associated markers that as such can serve as markers for T cell activation^3^. We indeed observed a slight upregulation of FOXP3, CTLA4 and TIGIT at mRNA and protein level in SF non-Treg but not near the levels observed in SF Treg (Fig. 1d, e and Supplementary Fig. 1b and 2b), further confirming that SF Treg and non-Treg are distinct cell populations. Together these data demonstrate that the inflammatory environment reinforces the Treg-associated program.

### Inflammatory environment derived Treg display a specific effector profile

A pairwise comparison between SF- and PB-derived Treg from healthy children revealed many differentially expressed genes, including core Treg markers including *FOXP3* and *CTLA4*, but also markers that reflect more differentiated Treg, like *PRDM1* (encoding Blimp-1), *ICOS, BATF* and *BACH2* (Fig. 2a, Supplementary Table 1). Based on recent literature we analyzed the expression of markers related to eTreg cell differentiation in mice. Hierarchical clustering analysis confirmed that Treg clustered separately from non-Treg and revealed a cluster of core Treg genes with increased expression in all Treg groups including *CTLA4*, *FOXP3*, *IL2RA* (encoding CD25), *TIGIT* and *IKZF4* (encoding Helios) (Fig. 2b, box 1). Furthermore, clustering revealed genes that show molecular heterogeneity within the Treg groups. Some markers were expressed almost exclusively by SF Treg (including *GMZB*, *ICOS* and *IL10*) whereas others were highly expressed by SF Treg but with shared, albeit lower expression by SF non-Treg (including *PRDM1*, *BATF*, *TNFRSF18* encoding GITR, *TNFRSF1B* and *HAVCR2* encoding TIM-3) (Fig. 2b, box 2). A third cluster concerns genes most highly expressed in non-Treg (Fig. 2b, box 3). Some genes in this cluster shared expression with SF Treg, such as *LAG3*, *CXCR3* and *PDCD1* (encoding PD-1) whereas expression of *CD40LG*, *BACH2*, *SATB1*, *CCR7* and *TCF7* was clearly lower in SF Treg (Fig. 2b, box 2 and 3). Interestingly, the more heterogeneously expressed genes listed in box 2 and 3 were previously associated with an eTreg profile, including *ICOS*, *IL10*, *PRDM1*, *TNRFSF18* and *PDCD1*. The increased expression of ICOS, GITR and PD-1 was verified at protein level (Fig. 2c and Supplementary Fig. 3a). Besides the differential expression of these markers, we found a significant enrichment of genes within SF Treg recently described in mice to be up- or downregulated in eTreg^20^ (Fig. 2d and Supplementary Fig. 3b), confirming the eTreg profile. Collectively, these data demonstrate that autoimmune-inflammation derived human Treg display an eTreg signature.

**Fig. 2.**
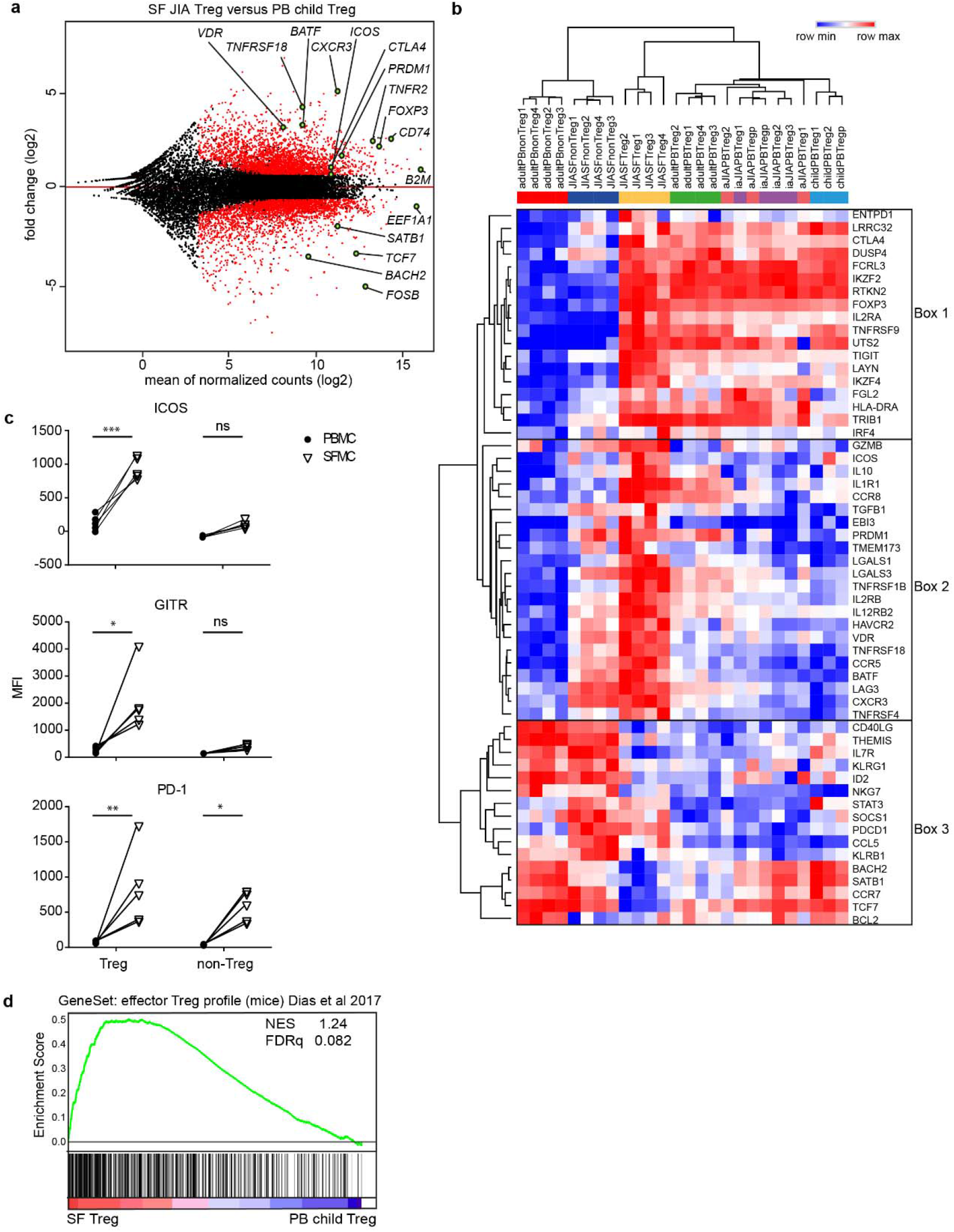
Inflammatory environment-derived Treg demonstrate a specific effector profile. **a** MA plot of the differentially expressed genes between synovial fluid (SF) and peripheral blood (PB) Treg of healthy children with black dots reflecting no change, and red dots transcripts with an adjusted p-value < 0.05, minimal mean of all normalized counts > 10 and log2 fold change > 0.6. **b** Heatmap with hierarchical clustering analysis including all groups measured with RNA-sequencing selected genes based on recent literature (relative expression of log2 RPKM). **c** MFI of ICOS, GITR and PD-1 in gated CD3^+^CD4^+^CD25^+^CD127^low^ Treg and CD3^+^CD4^+^CD25^−^ CD127^+^ non-Treg from paired SFMC and PBMC from 5 JIA patients. Statistical comparisons were performed using two-way ANOVA with Sidak correction for multiple testing: ns, * *P* < 0.05, \*\**P* < 0.01, \*\*\**P* < 0.001. Data are representative of two or more independent experiments. **d** GeneSet Enrichment Analysis (GSEA) of effector Treg genes in mice (identified by Dias *et al.*^29^) in pairwise comparisons involving SF Treg and PB Treg derived from healthy children, represented by the normalized enrichment score (NES) and FDR statistical value (FDRq).

### SF Treg demonstrate adaptation to the interferon-skewed inflammatory environment

To determine the relationship between inflammatory environment-derived and PB-derived cells, we performed unsupervised PCA analysis on SF Treg and non-Treg, and PB Treg from healthy children. PB Treg were clearly separated from SF-derived cells, confirming that environment plays a dominant role in determining the transcriptional landscape (Fig. 3a). K-mean clustering analysis showed that SF Treg and non-Treg share increased expression of genes related to inflammation-associated pathways, including pathways linked to cytokine responses such as Interferon and IL-12 (Supplementary Fig. 4a). K-mean, and subsequent gene ontology analysis of SF Treg and all PB Treg groups derived from children demonstrated that pathways associated with Th1-skewing were specifically upregulated in SF Treg (Fig. 3b and Supplementary Fig. 4b, c). Indeed, expression of the Th1 key transcription factor *TBX21* (T-bet), the Th1 related chemokine receptor *CXCR3*, and IL-12 receptorβ2 (*IL12RB2*) in SF Treg was increased on both mRNA and protein level (Fig. 3c, d and Supplementary Fig. 4d). Expression of *TBX21* and *CXCR3* was equally high in both SF Treg and non-Treg, whilst *IL12RB2* showed a significantly higher expression in SF Treg (adjusted p-value = 6.8E^−10^). Accordingly, SF Treg in contrast to non-Treg showed co-expression of T-bet and FOXP3 protein excluding significant contamination of non-Treg potentially contributing to the high T-bet levels observed in SF Treg (Fig. 3e). These results indicate that SF Treg exhibit functional specialization to allow regulation of specific Th cell responses at particular tissue sites, as was previously established in mice^21–23^. In line herewith, we found an enrichment in SF Treg of the transcriptional signature of TIGIT^+^ Treg which have been recently identified as activated Treg selectively suppressing Th1 and Th17 cells in mice^19^ (Supplementary Fig. 4e).

**Fig. 3.**
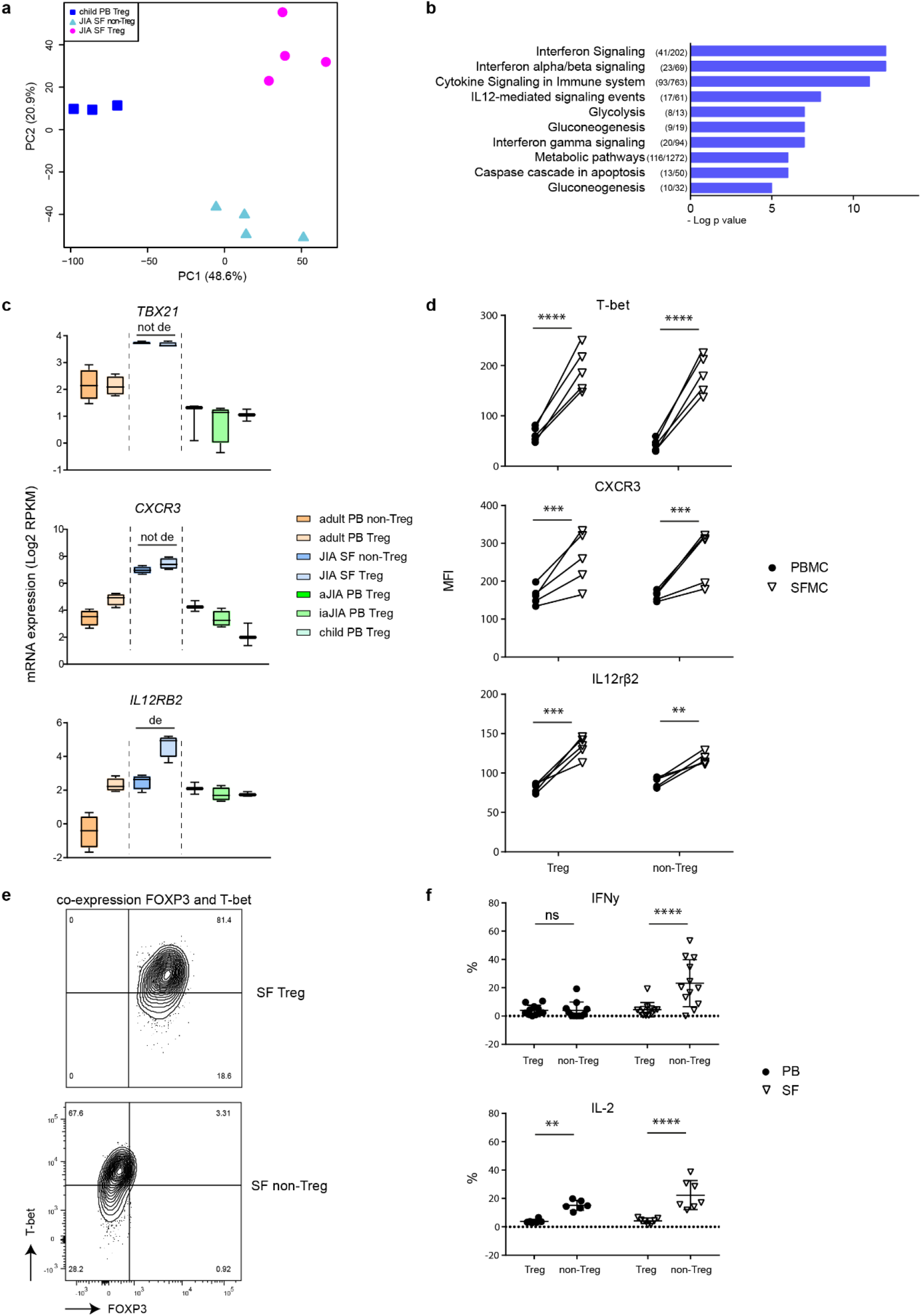
SF Treg adapt to the interferon-skewed inflammatory environment but lack cytokine production. **a** Unsupervised Principal Component Analysis (PCA) of synovial fluid (SF)-derived Treg and non-Treg and peripheral blood (PB)-derived Treg from healthy children. **b** Gene ontology terms related to the 1791 genes specifically upregulated in SF compared to PB Treg derived from children (active juvenile idiopathic arthritis (JIA), inactive JIA and healthy children), ranked by enrichment scores. The number of upregulated genes compared to the total number annotated in the gene ontology term are depicted before the terms. **c** mRNA expression (log2 RPKM) of *TBX21*, *CXCR3* and *IL12RB2* in Treg derived from PB of healthy adults (adult), healthy children (child), JIA patients with active (aJIA) or inactive (iaJIA) disease, SF of JIA patients (JIA SF) and non-Treg from PB of healthy adults and SF of JIA patients (de = differentially expressed according to description in Fig. 1d). **d** MFI of T-bet, CXCR3 and IL12Rβ2 in CD3^+^CD4^+^CD25^+^CD127^low^ Treg and CD3^+^CD4^+^CD25^−^CD127^+^ non-Treg from paired SFMC and PBMC from 5 JIA patients. **e** Representative contourplot of T-bet and FOXP3 in CD3^+^CD4^+^CD25^+^FOXP3^+^ Treg and CD3^+^CD4^+^CD25^int/−^FOXP3^−^ non-Treg from SFMC. **f** Percentage of IFNγ and IL-2 measured in CD3^+^CD4^+^CD25^+^FOXP3^+^ Treg and CD3^+^CD4^+^CD25^int/−^FOXP3^−^ non-Treg from SFMC and PMBC (IFNγ: n = 11 PB, n = 12 SF; IL-2: n = 6 PB, n = 7 SF). (**d** and **f**) Data are representative of two independent experiments. Statistical comparisons were performed using two-way ANOVA with Sidak correction for multiple testing. ns, \**P* < 0.05, \*\**P* < 0.01, \*\*\*\**P* < 0.0001.

The high expression of Th1-related proteins raised the question whether SF Treg may have acquired a Th1 phenotype. However, SF Treg failed to produce both IL-2 and IFNγ (Fig. 3f and Supplementary Fig. 4f) and responded dose-dependently to IL-2 with increasing pSTAT5 levels (Supplementary Fig. 4g), a signaling pathway pivotal for Treg survival and function^24^. In fact, compared to PB Treg, SF Treg appeared to be even more responsive to IL-2. Altogether, our findings demonstrate adaptation of SF Treg to their inflammatory environment while maintaining Treg key features.

### Effector Treg differentiation and inflammation are regulated by the (super-)enhancer landscape

To explore the mechanistic regulation of the inflammation-adapted eTreg profile we analyzed the enhancer landscape of SF Treg. Enhancers are distal regulatory elements in the DNA that allow binding of transcription factors and as such coordinate gene expression. Epigenetic regulation of enhancers is critical for context-specific gene regulation^25^. Enhancers can be defined by areas enriched for monomethylation of lysine 4 on histone H3 (H3K4me1 enrichment) and acetylation of lysine 27 on histone H3 (H3K27ac enrichment), where H3K27Ac identifies active enhancers^26^. ChIP-seq performed for H3K27ac and H3K4me1 using SF Treg and healthy adult PB Treg showed a striking difference in enhancer profile and activity, with 3333 out of 37307 and 970 out of 10991 different peaks called between SF and PB Treg for H3K27ac and H3K4me1, respectively (Fig. 4a, Supplementary Fig. 5b, upper panel and Supplementary Table 2). We next assessed if the transcriptome of SF Treg is reflected at an epigenetic level. Genes that demonstrated increased H3K27ac and/or H3K4me1 were increased at the mRNA level and *vice versa* (Fig. 4a, Supplementary Fig. 5b, middle and lowest panel and Supplementary Table 2), confirming that gene expression and chromatin-acetylation and -monomethylation are interconnected in these cells.

**Fig. 4.**
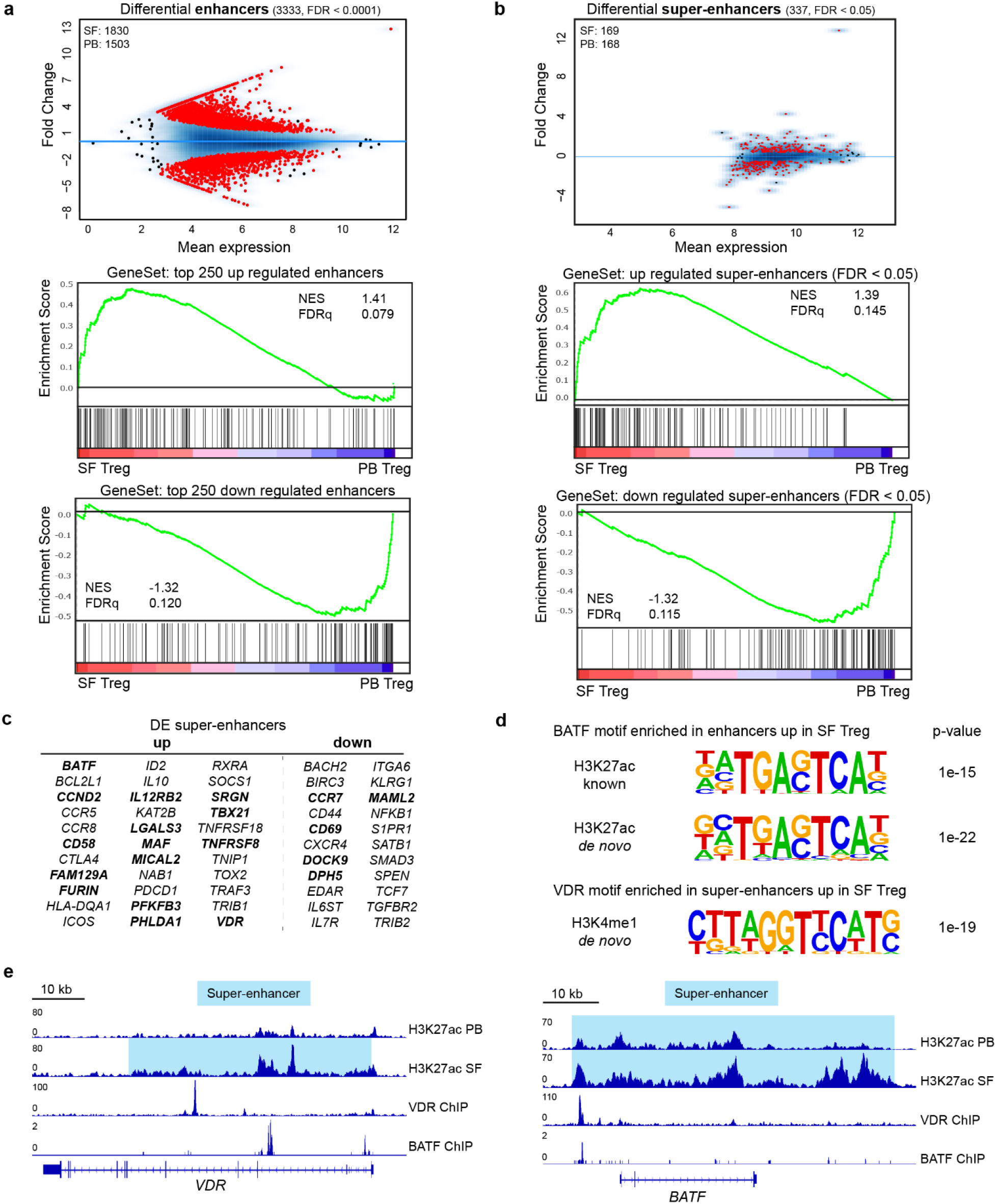
Environment-specific effector Treg profile is regulated by the (super-)enhancer landscape. **a** MA plots of differentially expressed enhancers (FDR<0.0001) in synovial fluid (SF) versus peripheral blood (PB) Treg for H3K27ac ChIP-seq with the number of SF- and PB-specific enhancers indicated (top). GeneSet Enrichment Analysis (GSEA) of the top 250 upregulated (middle) and downregulated (bottom) enhancers in pairwise comparisons involving transcriptome data of SF Treg and healthy adult PB Treg, represented by the normalized enrichment score (NES) and FDR statistical value (FDRq). **b** Same as in **a** but for super-enhancers (FDR < 0.05). **c** Selection of super-enhancers up- and downregulated in SF Treg versus healthy adult PB Treg for H3K27ac and H3K4me1 ChIP-seq (FDR < 0.05; bold = up/down for both and differentially expressed in SF versus PB Treg on transcriptome level, see also Supplementary Table 2). **d** Motifs, known and *de novo*, for transcription factor binding sites predicted using HOMER, enriched in the upregulated (super-)enhancers in SF Treg compared to healthy adult PB Treg for H3K27ac and H3K4me1 ChIP-seq. **e** Gene tracks for *VDR* and *BATF* (H3K27ac) displaying ChIP-seq signals, with the super-enhancer region highlighted in blue, in healthy adult PB Treg, SF Treg, VDR-specific (GSE89431) and a BATF-specific (GSE32465) ChIP-seq.

Super-enhancers are large clusters of enhancers that specifically regulate genes defining cell identity, both in health and disease^27,28^. Also at super-enhancer level, we found significant enrichment in the transcribed genes, again indicating that the SF Treg profile is mediated by epigenetic changes (Fig. 4b and Supplementary Fig. 5c, middle and lowest panel). The analysis of differential super-enhancer-associated genes revealed 337 out of 791 and 317 out of 713 different gene loci for H3K27ac and H3K4me1, respectively (Fig. 4b and Supplementary Fig. 5c, upper panel). Specifically, we identified that super-enhancers associated with genes related to canonical Th1 differentiation, such as *TBX21* and *IL12RB2*, are increased in SF Treg (Fig. 4c) demonstrating epigenetic regulation of the environment-associated profile. Core Treg genes encoding the effector molecules *ICOS*, *IL10, CTLA4*, *TNFRSF18* and *PDCD1* were associated with increased super-enhancers as well as *FURIN* and *ID2* which are more putative functional Treg markers. Importantly, eTreg differentiation was also reflected in the enhancer profile of SF Treg, with differential expression of super-enhancer-associated transcriptional regulators including *BATF*, *BACH2*, *SATB1, TRAF3* and *SOCS1*. Moreover, we found super-enhancers associated with markers not previously related to (e)Treg differentiation and mostly specific for human Treg, including *VDR*, *RXRA* (encoding Retinoic acid receptor RXR-alpha), *KAT2B* (encoding p300/CBP-associated factor (PCAF)), *TOX2* (encoding TOX High Mobility Group Box Family Member 2) and *IL12RB2*. Inflammation related homing markers (*CCR2*, *CCR5*, *CCR7*) were associated with differential H3K27ac levels in SF Treg as well. Finally, regulation of cell cycling and apoptosis was also mirrored in the super-enhancer landscape of SF compared to PB Treg by increased activity of *DPH5*, *MICAL2, PHLDA1* and *CCND2.* Altogether, these super-enhancer-associated genes reflect adaptation and specialization of Treg within an inflammatory environment.

Motif analysis for *in silico* prediction of transcription factor binding sites in (super-)enhancers specifically upregulated in SF compared to PB Treg revealed motifs for Treg-specific transcription factors. These included STAT5 and Myb; the latter recently described as a core transcription factor in eTreg differentiation^29^. Recent papers further demonstrated BATF and RelA as crucial regulators for eTreg in mice^30,31^. In support hereof, we found significant enrichment of BATF, TF65 (encoding RelA), and VDR motifs within the (super-)enhancer regions specifically upregulated in SF Treg (Fig. 4d and Supplementary Fig. 5d), thereby further strengthening the eTreg profile of SF Treg. The most prevalent motifs belong to the activator protein 1 (AP-1) transcription factor subfamily, part of the basic leucine zipper (bZIP) family (Supplementary Fig. 5d) indicating this subfamily is crucial in driving the eTreg signature, as similarly proposed by DiSpirito *et al.*^32^ for pan-tissue Treg. Moreover, within the super-enhancers of both the transcriptional regulators VDR and BATF the binding sites for each of them were found, indicating that BATF (in complex with other proteins) and VDR regulate their own gene expression, and also control other eTreg genes (Fig. 4e). Together, these observations demonstrate that the inflammation-adapted effector phenotype of SF Treg is regulated at an epigenetic level, with a super-enhancer profile and enrichment of binding sites for eTreg genes.

### The vitamin D3 receptor is a predicted key regulator of effector Treg differentiation

To extract key gene regulators in the differentiation from healthy PB to SF Treg a data-driven network and enrichment approach was performed (RegEnrich). The top predicted regulators driving the differentiation from PB to SF Treg were found to be *TCF7*, *LEF1*, *JUN*, *SPEN* (negative) and *ENO1*, *THRAP3* and *VDR* (positive) (Fig. 5a, b). *SPEN*, *THRAP3* and *VDR* have not been previously associated with eTreg differentiation. *VDR* associated downstream with core Treg genes including *FOXP3* and *STAT3*, but also *Tbx21* (Fig. 5a, c and Supplementary Table 3). In line herewith, there was a strong positive correlation for both FOXP3 (r=0.772, p <0.0001) and T-bet (r=0.937, p <0.0001) with VDR protein expression (Fig. 5e) reinforcing the predicted regulator network.

**Fig. 5.**
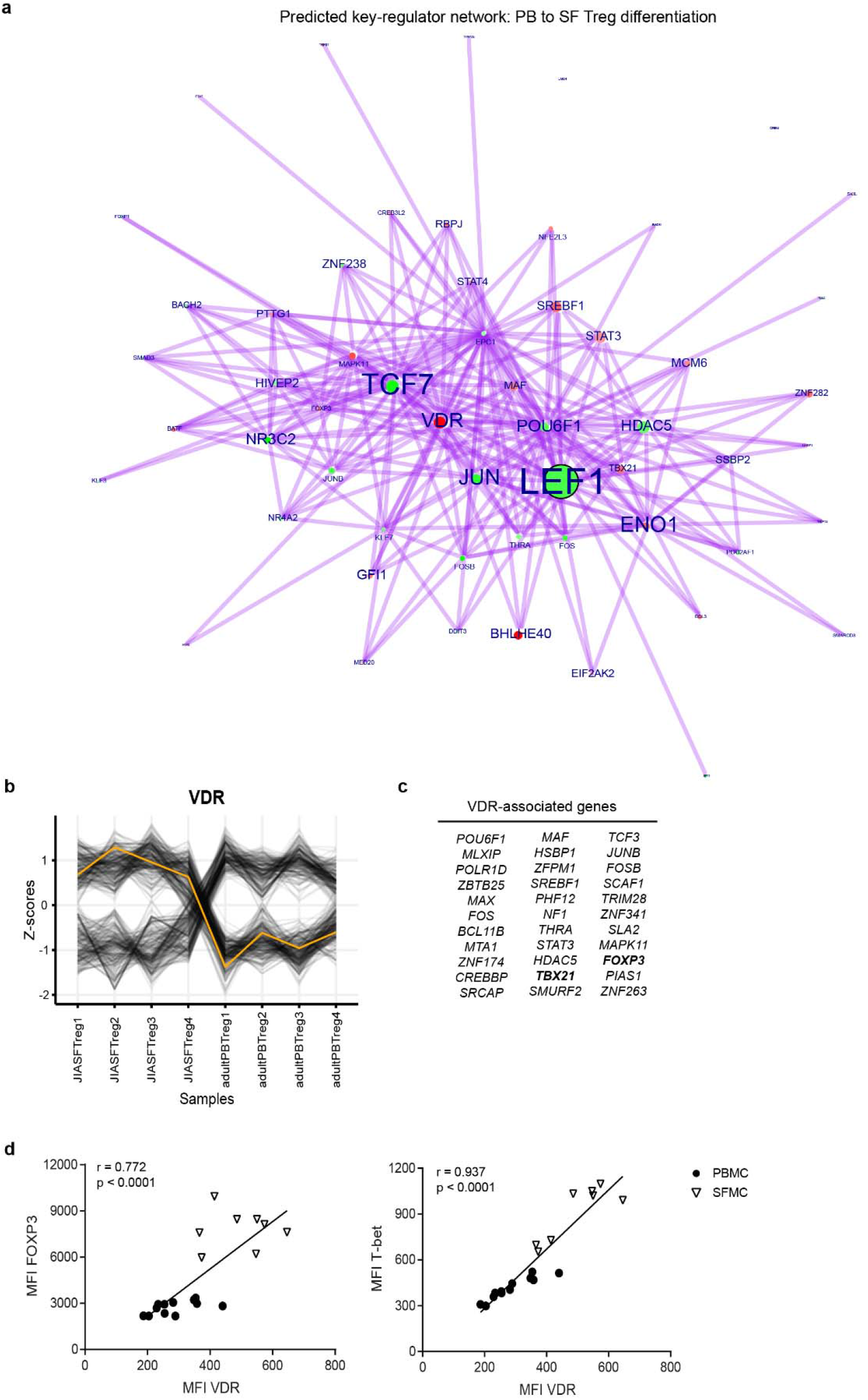
The vitamin D receptor is a predicted key-regulator of effector Treg differentiation. **a** Network inference of key-regulators driving peripheral blood (PB) to synovial fluid (SF) Treg differentiation on RNA level, based on unsupervised weighted correlation network analysis followed by Fisher’s exact test (red = upregulation, green = downregulation). Circle size indicates −log10(p) for each comparison, with the p(-value) derived from differential expression analysis (log fold change > 1), and the text size represents the RegEnrich score; for both, larger indicates higher scores (see also Supplementary Table 3). **b** The RNA expression profile of *VDR* from JIA SF Treg samples 1-4 to adult PB Treg samples 1-4 indicated by the yellow line, with its associated genes in grey; values are normalized to the Z-score. **c** All genes associated with VDR defined by the key-regulator network inference. **d** Pearson’s correlation plot of VDR and FOXP3 (left) and T-bet (right) MFI’s in PB (•) and SF (▼) with the line fitted by linear regression (n = 12 PB HC and PB JIA (PBMC), n = 8 SF JIA (SFMC)). Data are representative of two independent experiments.

### The effector Treg profile overlaps with other chronic inflammatory diseases and the human tumor Treg signature

To investigate whether the program identified in SF Treg is universal for human Treg exposed to inflammation, we compared our findings with Treg derived from PB and inflammatory joints of RA patients. Indeed, also in Treg from inflammatory exudate of RA patients similar eTreg genes were upregulated (Fig. 6a).

**Fig. 6.**
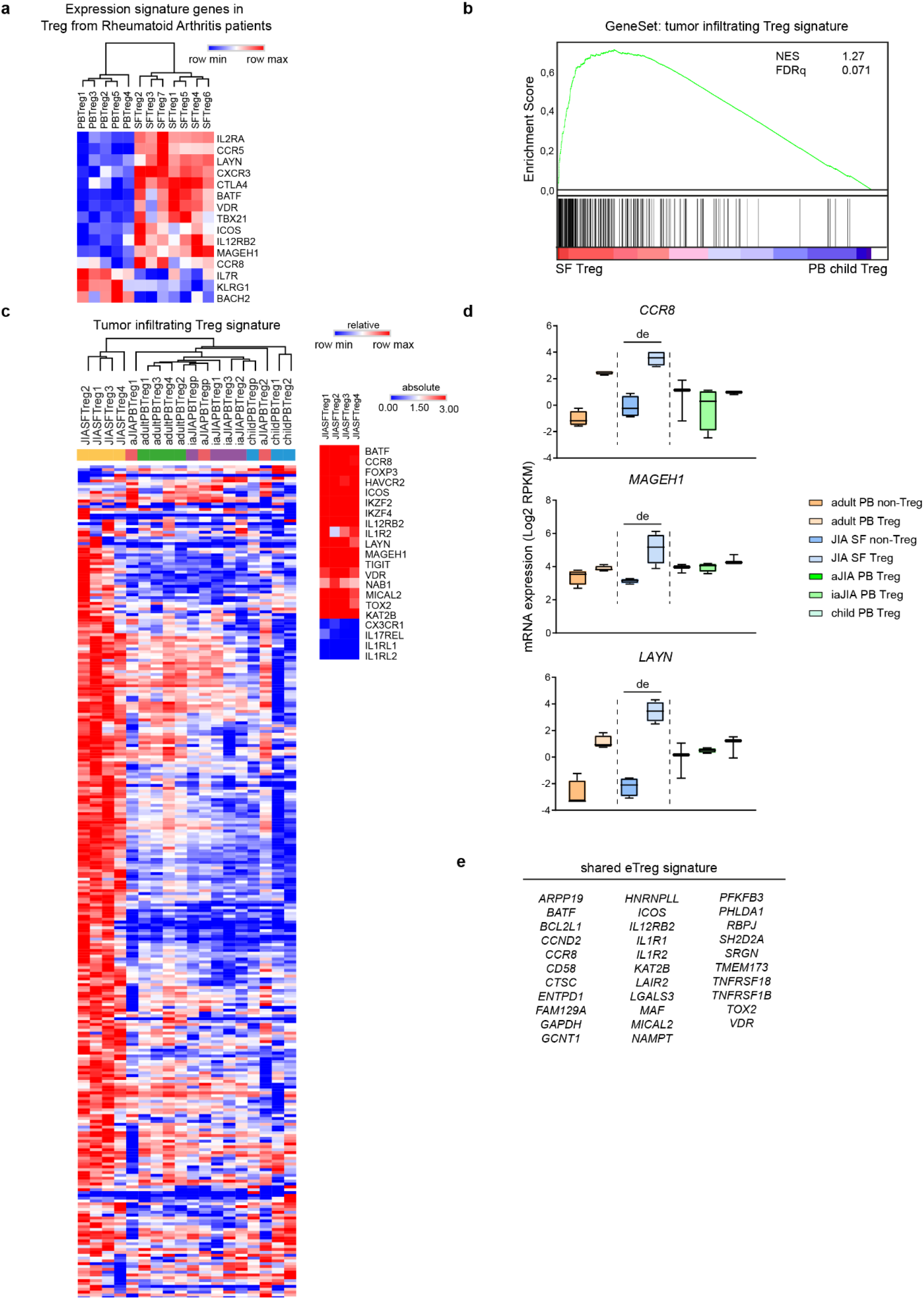
The effector Treg program is universal and overlaps with the human tumor-infiltrating Treg signature. **a** Heatmap with unsupervised hierarchical clustering analysis on peripheral blood (PB) and (partially paired) synovial fluid (SF) Treg from Rheumatoid Arthritis patients measured with a gene array, on selected signature genes identified from juvenile idiopathic arthritis (JIA) SF Treg and tumor-infiltrating Treg. **b** GeneSet Enrichment Analysis (GSEA) of tumor-infiltrating Treg signature genes (identified by De Simone *et al.*^13^) in pairwise comparisons involving SF Treg and healthy child PB Treg, represented by the normalized enrichment score (NES) and FDR statistical value (FDRq). **c** Heatmap with hierarchical clustering analysis including all groups measured with RNA-sequencing on the identified tumor-infiltrating Treg signature genes (ref. ^13^), with relative expression of log2 RPKM. Small heatmap on the right shows log2 RPKM values of a selection of genes in SF Treg. **d** mRNA expression (log2 RPKM) of *CCR8*, *MAGEH1* and *LAYN* in Treg from PB of healthy adults (adult), healthy children (child), JIA patients with active (aJIA) or inactive (iaJIA) disease, SF of JIA patients (JIA SF) and non-Treg from PB of healthy adults and SF of JIA patients as determined by RNA-sequencing analysis (de = differentially expressed according to description in Fig. 1d; adjusted p-values *CCR8* = 2.2E^−13^, *MAGEH1* = 1.2E^−07^, *LAYN* = 1.2E^−33^). **e** Identification of a human effector Treg profile based on the overlapping genes upregulated in JIA SF Treg, RA SF Treg and tumor-infiltrating Treg (identified in several studies, see refs. ^13–15,69,70^), and reflected on the (super-)enhancer landscape of SF Treg.

We then compared our data to recently published human tumor-infiltrating Treg specific signatures^13–15^. Strikingly, the tumor-infiltrating Treg signature from De Simone *et al.* was enriched in SF Treg (Normalized Enrichment Score (NES) 1.27, FDR statistical value (FDRq) 0.071; Fig. 6b). *Vice versa*, the differentially expressed genes in SF Treg versus PB Treg were enriched in tumor-infiltrating compared to normal tissue Treg (Plitas *et al.*, 2016: NES 1.69, FDRq 0.017, Magnuson *et al.,* 2018: NES 1.88, FDRq 0.002; Supplementary Fig. 6a, b). Hierarchical clustering analysis of the tumor-infiltrating gene signature from De Simone *et al.* further revealed separate clustering of SF Treg (Fig. 6c), indicating that the high expression of the signature genes is specific for human Treg at a site that is characterized with infiltration of immune cells. We also observed a small set of genes that were not upregulated in SF Treg, e.g. *CX3CR1*, *IL17REL*, *IL1RL1* and *IL1RL2*, implying environment-restricted adaptation (Fig. 6c). The three genes that were described as the most enriched and distinctive genes in tumor-infiltrating Treg, *LAYN*, *MAGEH1,* and *CCR8,* were selectively and highly upregulated in SF Treg (Fig. 6d). These data demonstrate that the Treg profile we observed is not restricted to the SF exudate from JIA and RA patients. In fact, it likely represents a more global profile of human Treg in inflammatory settings, likely fine-tuned by environment specific adaptations.

To investigate which cues might be specific for SF we explored the differences between upregulated genes in SF and tumor-infiltrating Treg. Relatively few pronounced SF-specific genes were found. On gene level, cytotoxic markers including *GZMM*, *GZMA* and *GNLY,* related to effector cell cytolysis and self-induced apoptosis of Treg^33^ were SF-specific (Supplementary Fig. 6c). Gene ontology biological process analysis further revealed that in SF both active cell cycling and apoptotic pathways were upregulated, whereas in the tumor microenvironment cell activation and effector responses were prominent (Supplementary Fig. 6d). Additionally, tumor-infiltrating Treg showed distinct expression of chemokines and chemokine receptors, e.g. IL-8 (CXCL8), CX3CR1, CXCR7 and CCR10 (Supplementary Fig. 6c), some of which have been proposed as prognostic marker and/or therapeutic target in cancer^34^.

From the JIA, RA and tumor datasets we could deduce a set of genes that appear universally upregulated in eTreg, and also being reflected in the (super-)enhancer landscape of SF Treg (Fig. 6e). *ICOS*, *BATF, MAF*, *TNFRSF18* and *SRGN* were revealed as core markers previously also related to mouse eTreg. Additionally, *IL12RB2, VDR,* and *KAT2B,* were identified as core genes, as well as *PFKFB3* and *LDHA* involved in glycolysis and the apoptosis-related genes *MICAL2* and *TOX2.* The latter two have so far not been linked to Treg in human or mice, although both have independently been linked to human cancers^35,36^. Collectively, our results demonstrate that human Treg in different inflammatory settings share an effector profile and strongly indicates that inflammation drives eTreg differentiation in a comparable manner in different environments.

## Discussion

The discovery of eTreg in non-lymphoid tissues and inflammatory sites in mice have raised questions how human eTreg are transcriptionally and epigenetically programmed in different settings. The present study provides evidence that human Treg in a local inflammatory setting undergo further differentiation into specialized eTreg, with a unique expression profile. This is reflected by high expression of TBX21 and CXCR3 and core Treg markers including FOXP3, CTLA4 and TIGIT. Essential Treg features are maintained, such as suppressive function, IL-2 responsiveness and absence of cytokine production. The transcriptional changes were mirrored at an epigenetic level, both in the enhancer and super-enhancer landscape, demonstrating that the profile is highly regulated. Additionally, we identify previously unappreciated and specific markers in human eTreg, like VDR, IL12Rβ2, and BATF; with VDR as a predicted key-regulator in human eTreg differentiation. Finally, we demonstrate that the profile is not limited to autoimmune inflammation in JIA and RA, but shares almost complete overlap with tumor-infiltrating Treg. The similarity between a tumor and an autoimmune setting might seem counterintuitive, since the former reflects an immune-suppressive environment whereas the latter is associated with immune activation. Both environments however share features including immune cell infiltration and inflammation^37^. This overlapping profile might therefore represent a more universal profile of human Treg in inflammatory environments. Because of the extensive genome-wide changes of Treg from an inflammatory environment insight in their functionality is important. In line with previous reports^16,17^, we show that SF Treg are indeed suppressive and additionally highly responsive to IL-2 by phosphorylation of STAT5, excluding impaired IL-2 signaling^38,39^. This advocates against an intrinsic defect of Treg in inflammation. Maintenance of inflammation may be explained by resistance of local effector cells, previously demonstrated in both JIA and RA^17,40,41^.

In mice, functional specialization of Treg towards Th1 inflammation by upregulation of T-bet is described as essential to prevent Th1 autoimmunity^9,10^. The remarkable co-expression of T-bet and FOXP3 in SF Treg in our study, with T-bet levels as high as in SF non-Treg, implies that functional specialization is translatable to a human setting. The increased expression of migration markers such as CXCR3 and CCR5 on protein, transcriptional and enhancer level further strengthens this, as high T-bet expression is crucial for migration and thus co-localization of Th1-specific Treg and Th1 effector T cells via upregulation of these chemokine receptors^9,10^. Contamination of Treg with non-Treg can be a concern when sorting *ex vivo* human cells. However, we exclude substantial contamination because *1)* our data show robust transcriptomic differences between SF Treg and non-Treg, with increased expression of Treg hallmark genes in SF Treg, *2)* sorted Tregs are >90% FOXP3^+^, *3)* there is clear co-expression of FOXP3 and T-bet on the single cell level, *4)* we show that SF Treg, while expressing high T-bet levels, lack cytokine production, potentially caused by high expression of FOXP3, SOCS1, EGR2 and EGR3 (Supplementary Table 1)^42,43^. Nevertheless, as recently shown by Miragaia *et al.*^4^ there may be heterogeneity within the eTreg population.

Our study is the first to investigate both the transcriptional and epigenetic regulation of eTreg in human inflammatory settings allowing us to discern human-specific eTreg regulation as well as commonalities between mice and men. The epigenetic landscape of human Treg is little explored, with only two papers studying H3K27acetylation and H3K4monomethylation of circulating human Treg^44,45^. Arvey *et al.* conclude that the majority of Treg lineage-specific elements are not conserved between mice and human further emphasizing the lack of knowledge of the human Treg epigenetic landscape. Gao *et al.* focused on SNPs within enhancers of PB Treg from Type 1 diabetes patients, and in line with our data showed, that cell cycle and apoptosis regulation is highly reflected in the enhancer landscape of these patients. These data further underscores the need for additional studies, especially in non-lymphoid tissues during non-homeostatic conditions. We found similarities between mice and men in eTreg differentiation regarding downregulation of *SATB1*, *BACH2, TCF7* and *LEF1*, all associated with a super-enhancer in PB but not SF Treg. Although all four genes are crucial for Treg development, recent reports show that downregulation of these regulators is necessary for further differentiation of Treg and preventing conversion into Th cells^46–52^. BACH2 can transcriptionally repress *PRDM1* (encoding Blimp-1) in T cells^50^, and in line herewith we observed increased acetylation and gene expression of *PRDM1* in SF Treg. This was further supported by the decrease in *TCF7* and *LEF1,* both negatively correlating with eTreg marker expression including *PRDM1*^52^. Moreover, we found BATF as a prominent marker in SF Treg, with increased expression on both transcriptional and epigenetic level, and a binding site in upregulated super-enhancer associated genes. In addition, high BATF expression was also observed in RA SF Treg and tumor-infiltrating Treg, suggesting BATF is a key transcriptional regulator in both human and mice eTreg. We also found remarkable differences in eTreg signature markers compared to what is known from mice. Most pronounced is upregulation of IL12RB2 at both the transcriptional and epigenetic level, and a binding site in upregulated super-enhancer associated genes, whereas in mice eTreg this marker is downregulated. Instead, impaired IL-12Rβ2 expression has been reported as a key checkpoint preventing Treg from fully differentiating towards Th1 cells in mice^23^. However, our data are in line with recent reports showing selective expression of *IL12RB2* in human PB Treg^44,53^, suggesting this marker has different functions in human and mice.

Another unexpected finding is the upregulation of VDR in both JIA and RA SF Treg, on both transcriptional and epigenetic level, as well as in tumor-infiltrating Treg. Although not highlighted, high VDR levels were present in breast tumor-infiltrating Treg^14^ and uterine eTreg^54^. Besides the well-known tolerogenic effects of vitamin D_3_^55^, VDR is not well-studied in the Treg context, especially in humans. A recent study did show that memory CCR6^+^ Th cells gained a suppressive phenotype, including CTLA-4 expression, and functional suppressive capacity similar to Treg upon incubation with vitamin D_3_^56^. Here we show VDR is a predicted key-regulator in SF eTreg, positively correlated with both FOXP3 and T-bet expression.

Another striking difference is absence of *KLRG1* upregulation in human eTreg. In mice, KLRG1 is a key marker to identify eTreg, although not crucial for eTreg function^57^. KLRG1 is not upregulated on SF Treg and associated with a super-enhancer only in PB Treg. Additionally, in our data gene expression of KLRG1 is restricted to non-Treg (both SF and PB). Confusing the findings are discrepancies possibly caused by the tissue measured or comparison made. Miragaia *et al.*^4^ showed upregulation of *PIM1* in non-lymphoid Treg compared to PB Treg in mice, but *PIM1* was not expressed in human tissue Treg. In our data however, *PIM1* is highly increased in SF compared to PB Treg of healthy adults and children (Supplementary Table 1). In summary, we show that human eTreg show similarities in eTreg differentiation markers identified in mice, but also found human-specific eTreg markers including IL-12Rβ2 and VDR, and absence of KLRG1.

Previous research has shown that *ex vivo* human Treg are highly glycolytic^58^, in line with their high proliferative capacity^59^. In mice increased glycolysis is required for eTreg differentiation and migration to inflammatory sites^60,61^. In support hereof, glycolysis-associated genes including *PFKFB3, LDHA,* and *PKM2* were increased in SF Treg on both RNA and (super-)enhancer level. Additionally, preliminary analysis of T-cell glucose consumption by Seahorse technology showed increased glycolysis in SF compared to PB Treg (data not shown). Finally, *ENO1*, encoding the glycolytic enzyme Enolase 1, is a predicted key-regulator of eTreg differentiation. Lately, application of Treg-based therapies for autoimmune diseases and transplantation settings is gaining renewed interest. Promising data from animal models and clinical trials, both with cell therapy involving adoptive transfer and with low dose IL-2 administration, pave the way for Treg therapies to reach the clinic^62^. Our data show that circulating human Treg are markedly different from their counterparts derived from sites characterized by immune activation. This is reflected in the expression of effector markers and transcriptional regulators, but also in the expression of specific chemokine receptors. Regarding Treg-based therapies appreciating that specific environments may require adapted Treg, both for migration and function, is important. Moreover, in this study we extensively profiled human eTreg abundantly present at sites of inflammation. This information can form a basis for follow-up studies that may eventually allow characterization of small amounts of circulating eTreg, for example to monitor patients undergoing treatment.

In conclusion, our study is the first to uncover the transcriptional and epigenetic program that defines human inflammatory Treg. SF Treg display an environment-adapted as well as an eTreg phenotype established at the epigenetic level. Moreover, we describe striking similarities of the eTreg program with tumor-infiltrating Treg and SF Treg from RA patients. This revealed a set of genes shared with human eTreg from affected sites in JIA, RA and cancer including BATF, VDR, MICAL2, TOX2, KAT2B, PFKFB3 and IL12Rβ2. Finally, we show that THRAP3, ENO1 and VDR are predicted key-regulators driving human SF eTreg differentiation at the transcriptomic level.

## Methods

### Collection of SF and PB Samples

Patients with JIA (n=41) were enrolled by the Pediatric Rheumatology Department at University Medical Center of Utrecht (The Netherlands). Of the JIA patients n=8 were diagnosed with extended oligo JIA and n=33 with oligo JIA, according to the revised criteria for JIA^63^, with an average age of 11.3 years (range 3.2-19 years) and a disease duration at the time of inclusion of 4.9 years (range 0.1-15 years). Patients with RA (n=7) were enrolled by the rheumatology outpatient clinic at Guy’s and St. Thomas’ Hospital NHS Trust (United Kingdom). The average age of RA patients was 61 years (range 30-75 years). PB and synovial fluid (SF) was obtained when patients visit the outpatient clinic via vein puncture or intravenous drip, and by therapeutic joint aspiration of the affected joints, respectively. The study was conducted in accordance with the Institutional Review Board of the University Medical Center Utrecht (approval no. 11-499/C; JIA) and the Bromley Research Ethics Committee (approval no. 06/Q0705/20; RA). PB from healthy adult volunteers (HC, n=20, average age 41.7 years with range 27-62 years) was obtained from the Mini Donor Service at University Medical Center Utrecht. PB from n=8 healthy children (average age 11.4 years with range 7.3-15.6 years) was obtained from a cohort of control subjects for a case-control clinical study.

### Isolation of SFMC and PBMC

SF of JIA patients was incubated with hyaluronidase (Sigma-Aldrich) for 30 min at 37°C to break down hyaluronic acid. Synovial fluid mononuclear cells (SFMCs) and peripheral blood mononuclear cells (PBMCs) were isolated using Ficoll Isopaque density gradient centrifugation (GE Healthcare Bio-Sciences, AB) and were used after freezing in Fetal Calf Serum (FCS) (Invitrogen) containing 10% DMSO (Sigma-Aldrich).

### Suppression assay

CD3^+^CD4^+^CD25^+^CD127^low^ cells (Treg) were isolated from frozen PBMC, using the FACS Aria III (BD). Antibodies used for sorting are: anti-human CD3-BV510 (OKT3), CD25-PE/Cy7 (M-A251; BD), CD127-AF647 (HCD127; Biolegend), CD4-FITC (RPA-T4; eBioscience). To check for FOXP3 expression of the sorted populations cells were fixed and permeabilized by using eBioscience Fixation and Permeabilization buffers (Invitrogen) and stained with anti-human FOXP3-eF450 (PCH101; eBioscience). Read out of proliferation is performed with the following antibodies: CD3-PerCP/Cy5.5 (UCHT1; Biolegend), CD4-FITC (RPA-T4; eBioscience), CD8-APC (SK1; BD). Total PBMC were labeled with 2μM ctViolet (Thermo Fisher) and cultured alone or with different ratios of sorted Treg (1:16, 1:8, 1:4, 1:2). Cells were cultured in RPMI1640 media containing 10% human AB serum with addition of L-Glutamine and Penicillin/Streptomycin. PBMC were stimulated by 0,1 μg/ml coated anti-CD3 (eBioscience) and incubated for four days in a 96 well round bottom plate (Nunc) at 37°C. After 4 days cells were stained with CD3, CD4, and CD8 for read out of proliferation by flow cytometry performed on FACS Canto II (BD Biosciences) and data was analyzed using FlowJo Software (Tree Star Inc.).

### STAT5 Phosflow

PBMC and SFMC were thawed and resuspended in PBS (0,5 − 1,0 × 10^6^ living cells/tube). Surface staining of CD3 (Biolegend) and CD4 (eBioscience) was performed for 25 min at 4°C. Cells were then stimulated with 0, 1.0, 10 or 100 IU/ml human (h)IL-2 (Proleukin; Novartis) for 30 min at 37°C, fixated and permeabilized by using buffers from the Transcription Factor Phospho Buffer Set (BD Biosciences). Intranuclear staining of FOXP3-eF450 (PCH101), T-bet-eF660 (eBio4B10; eBioscience), CD25-PE/CY7 (M-A251) and pSTAT5-PE (pY695; BD) was performed for 50 min at 4°C. Data acquisition and analysis as above.

### RNA-sequencing

CD3^+^CD4^+^CD25^+^CD127^low^ and CD3^+^CD4^+^CD25^−^CD127^+^ cells were sorted by flow cytometry from HC PBMC and JIA patient SFMC and PBMC. Total RNA was extracted using the AllPrep DNA/RNA/miRNA Universal Kit (Qiagen) as specified by the manufacturer’s instructions and stored at −80°C. Sequencing libraries were prepared using the Rapid Directional RNA-Seq Kit (NEXTflex). Libraries were sequenced using the Nextseq500 platform (Illumina), producing single end reads of 75bp (Utrecht Sequencing Facility). Sequencing reads were mapped against the reference human genome (hg19, NCBI37) using BWA (v0.7.5a, mem ‒t 7 ‒c 100 ‒M -R) (see Supplementary table 4 for quality control information). Differential gene expression was performed using DEseq2 and custom Perl and R (www.r-project.org) scripts (https://github.com/UMCUGenetics/RNASeq). For K-means clustering and PCA analysis, genes with fold change between samples on 10th and 90th quantile at least 1 log2 RPKM and expression at least 2 log2 RPKM in the sample with the maximal expression were used. K-means clustering was done on gene expression medians per group, with an empirically chosen k of 14. Gene Ontology pathway analyses were performed using ToppFun (https://toppgene.cchmc.org/enrichment.jsp) with as input genes belonging to the defined k-mean clusters, with an FDR-corrected p-value <0.05 defining significance. Gene set enrichment analysis (GSEA)^64^, with as input the log2 RPKM data, was used to assess whether specific signatures were significantly enriched in one of the subsets. One thousand random permutations of the phenotypic subgroups were used to establish a null distribution of enrichment score against which a normalized enrichment score and FDR-corrected q values were calculated. Gene-sets were either obtained by analyzing raw data using GEO2R (NCBI tool) or downloaded from published papers, or self-made based on the H3K27ac/H3K4me1 data. In particular, the following published data sets were used: human core Treg signature: Ferraro *et al.*^65^; effector Treg signature in mice: Dias *et al.*^29^; tumor-infiltrating Treg signature: De Simone *et al.*^13^, Plitas *et al.*^14^, Magnuson *et al.*^15^; effector Treg genes in mice: gene set GEO: GSE61077; TIGIT^+^ Treg signature in mice: Joller *et al.*^19^. Identification of key-regulators was performed using RegEnrich (https://bitbucket.org/systemsimmunology/regenrich/src/master/) based on the differential gene expression data followed by unsupervised WGCNA for the network inference, mean RPKM counts > 0 were included, and Fisher’s exact test was used for enrichment analysis. Heatmaps and subsequent hierarchical clustering analyses using One minus Pearson correlation were performed using Morpheus software (https://software.broadinstitute.org/morpheus/).

### H3K27ac and H3K4me1 ChIP-sequencing

ChIP-seq was performed as described previously by Peeters *et al.*^66^. In short, PBMC from HC and SFMC from JIA patients were thawed and 0.5-1 million CD3^+^CD4^+^CD25^+^CD127^low^ cells were sorted by flow cytometry. For each sample, cells were crosslinked with 2% formaldehyde and crosslinking was stopped by adding 0.2 M glycine. Nuclei were isolated in 50 mM Tris (pH 7.5), 150 mM NaCl, 5 mM EDTA, 0.5% NP-40, and 1% Triton X-100 and lysed in 20 mM Tris (pH 7.5), 150mMNaCl, 2mMEDTA, 1% NP-40, 0.3% SDS. Lysates were sheared using Covaris microTUBE (duty cycle 20%, intensity 3, 200 cycles per burst, 60-s cycle time, eight cycles) and diluted in 20 mM Tris (pH 8.0), 150 mM NaCl, 2 mM EDTA, 1% X-100. Sheared DNA was incubated overnight with anti-histone H3 acetyl K27 antibody (ab4729; Abcam) or anti-histone H3 (mono methyl K4) antibody (ab8895; Abcam) pre-coupled to protein A/G magnetic beads. Cells were washed and crosslinking was reversed by adding 1% SDS, 100mM NaHCO3, 200mM NaCl, and 300 μg/ml proteinase K. DNA was purified using ChIP DNA Clean & Concentrator kit (Zymo Research), end-repair, a-tailing, and ligation of sequence adaptors was done using Truseq nano DNA sample preparation kit (Illumina). Samples were PCR amplified, checked for the proper size range and for the absence of adaptor dimers on a 2% agarose gel and barcoded libraries were sequenced 75 bp single-end on Illumina NextSeq500 sequencer (Utrecht DNA sequencing facility). Sample demultiplexing and read quality assessment was performed using BaseSpace (Illumina) software. Reads with quality score of Q>30 were used for downstream analysis. Reads were mapped to the reference genome (hg19/hg38) with Bowtie 2.1.0 using default settings for H3K27ac ChIP-seq and BWA for H3K4me1 ChIP-seq (see Supplementary Table 4 for quality control information). SAM files were converted to BAM files using samtools version 0.1.19. Peaks were subsequently called using MACS-2.1.0. Enriched regions were identified compared to the input control using MACS2 callpeak--nomodel --exttsize 300 --gsize=hs -p 1e-9. The mapped reads were extended by 300bp and converted to TDF files with igvtools-2.3.36 and were visualized with IGV-2.7.2^67^ for H3K27ac ChIP-seq and H3K4me1 ChIP-seq. Differential binding analysis was performed using the R package DiffBind v1.8.5. In DiffBind read normalization was performed using the TMM technique using reads mapped to peaks which were background subtracted using the input control. Enhancer gene associations were determined as the nearest TSS to the center of the enhancer and super-enhancer locus. Super-enhancers were identified by employing the ROSE algorithm^68^ using a stitching distance of the MACS2 called peaks of 12.5kb, peaks were excluded that were fully contained in the region spanning 1000bp upstream and downstream of an annotated TSS (-t 1000). The H3K27ac/H3K4me1 signal was corrected for background using the input control and subsequently ranked by increasing signal (Fig. S5A). Super-enhancer gene associations were determined as the nearest TSS to the center of the enhancer and super-enhancer locus. BEDtools v2.17.0 was used for general manipulation of peak bed-files. Motif enrichment analysis was performed using the HOMER software v4.11 (findMotifsGenome.pl; hg19/hg38; -size 200). ChIP-seq data for activated Treg, VDR and BATF was retrieved from GEO:GSE43119, GSE89431 and GSE32465, respectively.

### Vitamin D receptor and 1,25 vitamin D3 incubation assay

Thawed HC PBMC and JIA patient SFMC were stained by surface and intranuclear staining as described above with the following antibodies: anti-human fixable viability dye eF506, FOXP3-PerCP/Cy5.5 (PCH101; eBioscience), CD3-AF700 (UCHT1), CD4-BV785 (OKT4; Biolegend), CD25-BV711 (2A3), CD127-BV605 (A019D5; Sony Biotechnology), and VDR-PE (D-6; Santa Cruz Biotechnology). Data acquisition and analysis as above.

### Microarray RA Treg

CD14^+^ monocytes (purity >98%) were depleted through positive selection using CD14 MicroBeads (Miltenyi Biotec). CD4^+^ T cells were isolated from the CD14^−^ cell fraction by negative selection (Miltenyi Biotec) and stained with CD4-PerCP/Cy5.5 (SK3), CD45RA-APC/Cy7 (HI100), CD45RO-PB (UCHL1), CD127-FITC (A019D5; BioLegend), and CD25-PE (REA945; Miltenyi Biotec). CD4^+^CD45RA^−^CD45RO^+^CD25^+^CD127^low^ cells (memory Treg cells), were sorted using a BD FACSAria II. Sorted Treg cell samples were lysed in 1000 μl of TRIzol (Invitrogen). Chloroform (200 μl) was added, and the samples were then whirl mixed and incubated for 2–3 minutes at room temperature. Following centrifugation (10,000g for 15 minutes at 4°C), the water phase was further purified using the ReliaPrepTM RNA Miniprep system (Promega) per manufacturer’s instructions. RNA integrity was confirmed on an Agilent Technologies 2100 bioanalyzer. One hundred nanograms of total RNA was used to prepare the targets (Affymetrix) in accordance with the manufacturer’s instructions. Hybridization cocktails were hybridized onto a Human Gene 2.0 ST Array. Chips were scanned and gene expression data were normalized using the RMA algorithm. Gene expression analysis was performed using Qlucore Omics Explorer software, version 3.0.

### Statistical analysis

For ChIP-seq and RNA-seq analysis, p-values were adjusted with the Benjamini-Hochberg procedure. Protein and cytokine expression was analyzed with Pearson’s correlation, two-tailed Mann-Whitney test, Two-way ANOVA with Sidak correction for multiple testing or a mixed-effects model with Dunnett post-hoc on paired data with missing values using GraphPad Prism.

## Supporting information

Sub table 2

Sub table 3

Sub Table 4

## Data availability

The authors declare that all data supporting the findings of this study are available within the article and its Supplementary information files or from the corresponding author upon reasonable request. RNA-seq and ChIP-seq raw data is deposited in the Gene Expression Omnibus and will be made publicly available upon acceptance in a journal.

## Acknowledgments

We would like to thank J. van Velzen and P. Andriessen van der Burght for technical assistance. F. van Wijk is supported by a VIDI grant from ZonMw (91714332).

## Author contributions

Conceptualization: G.M. and L.L., Fv.W and J.v.L. Performed experiments: G.M., L.L. M.v.d.W., R.S., M.K., S.Ve., A.P. Data analysis: G.M., L.L., M.M., M.v.d.W., J.G.C.P., A.P., W.T., and J.v.L. Patients selection and clinical interpretation: G.M., S.Va., and S.d.R. Supplied RA dataset: V.F., C.R., and L.S.T. Writing: G.M., L.L., M.v.d.W., J.v.L., and F.v.W. Supervision: J.v.L. and F.v.W. All authors reviewed and edited the manuscript.

