## Supplementary figures and images for "Conserved human effector Treg signature is reflected in transcriptomic and epigenetic landscape"

### Fig S1

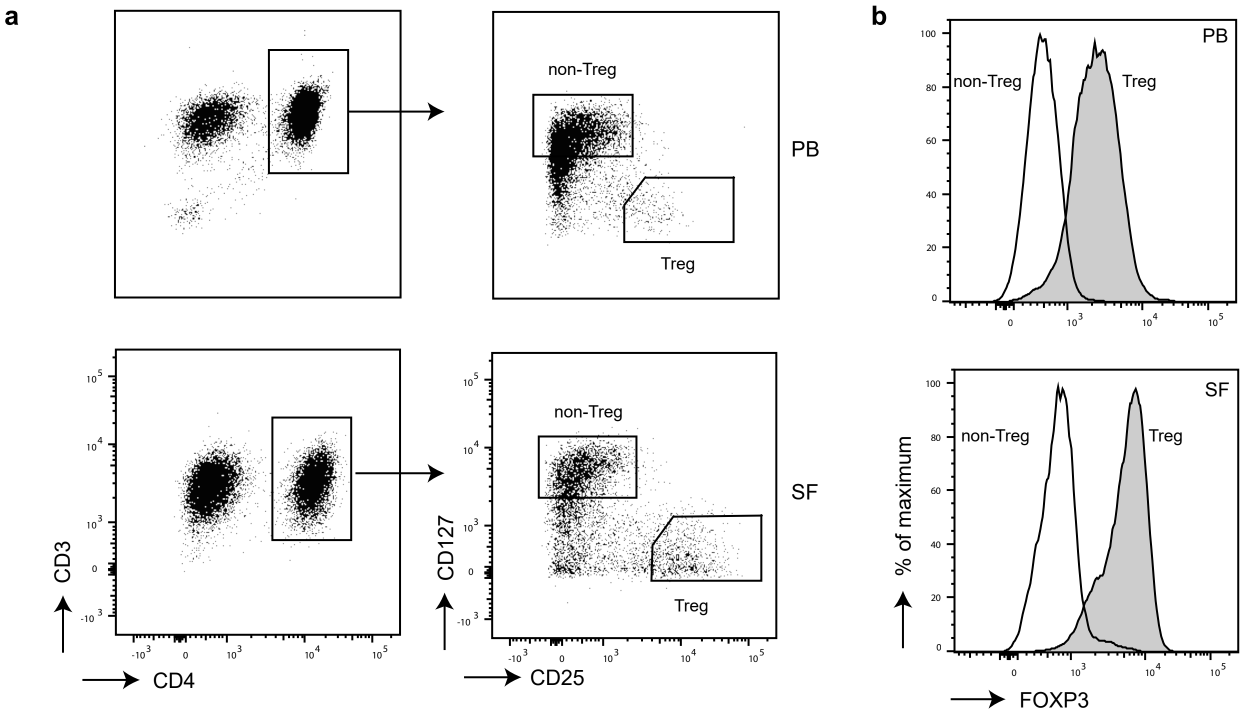

### Fig S2

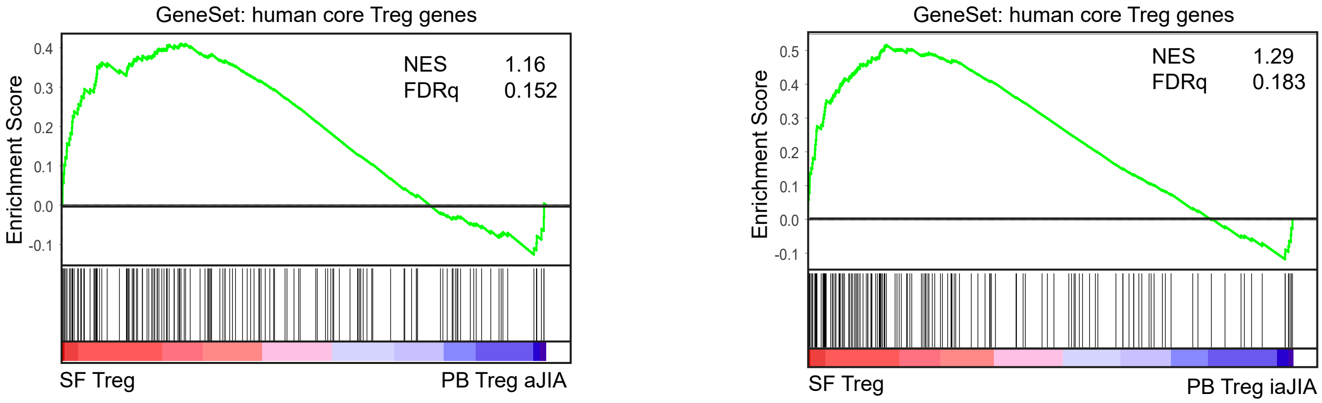

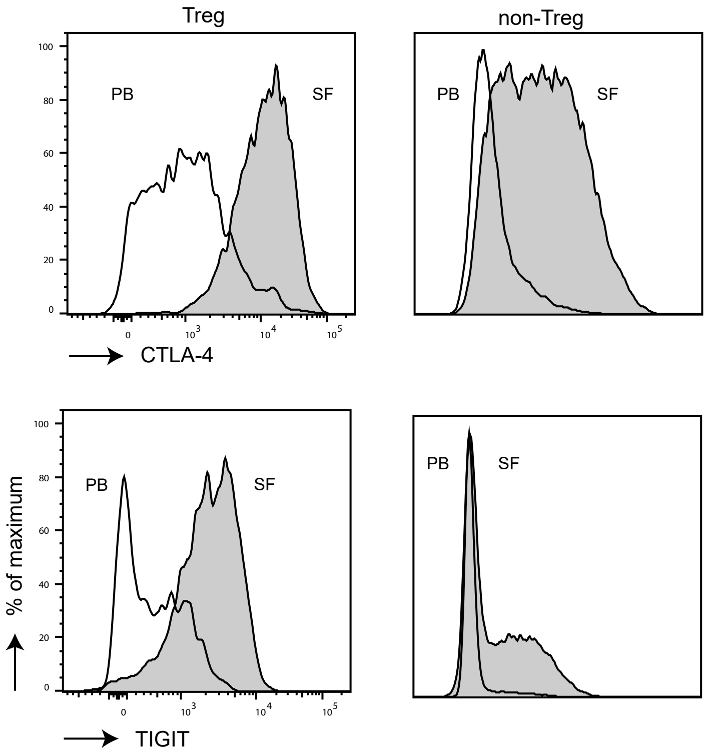

### Fig S3

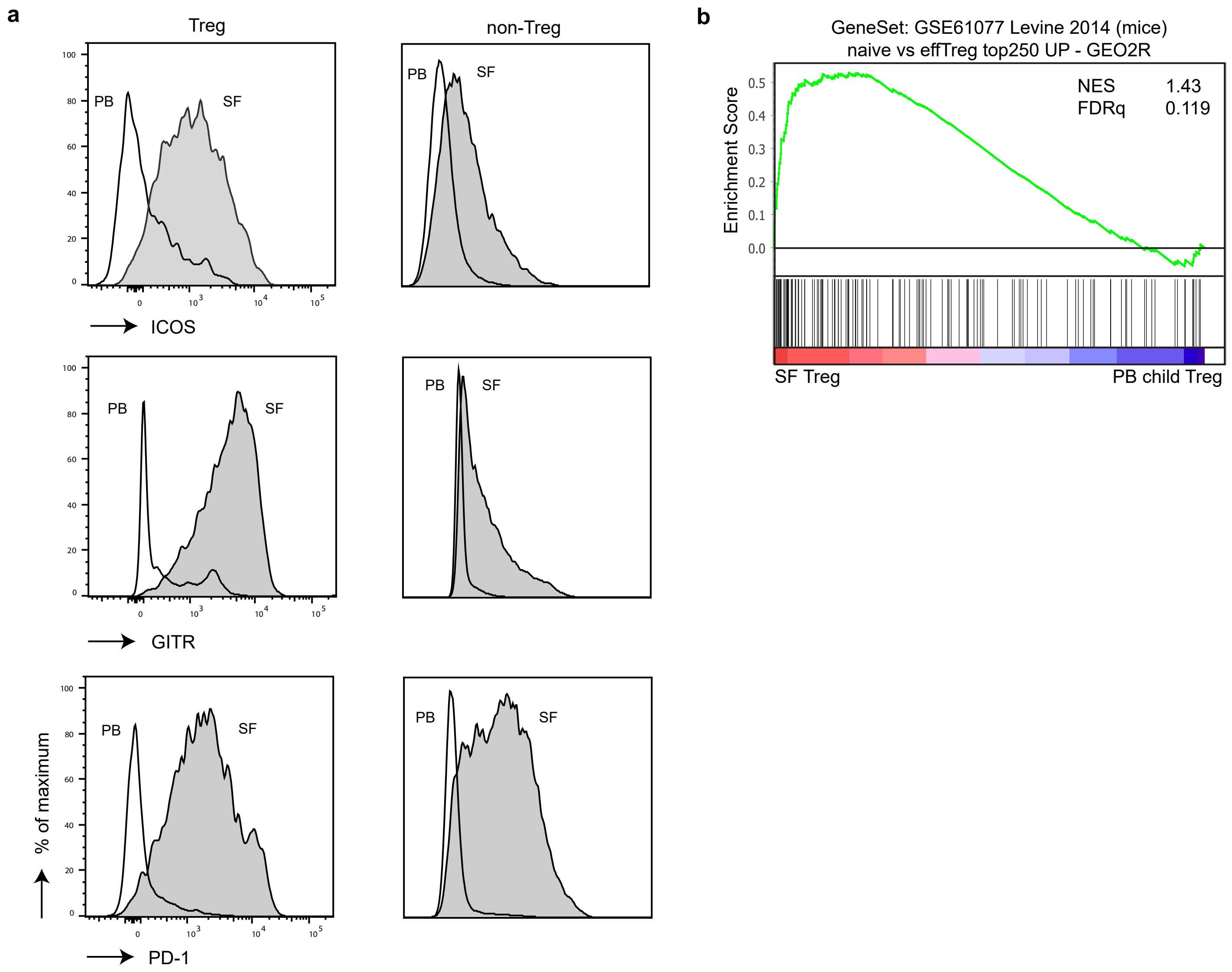

### Fig S4

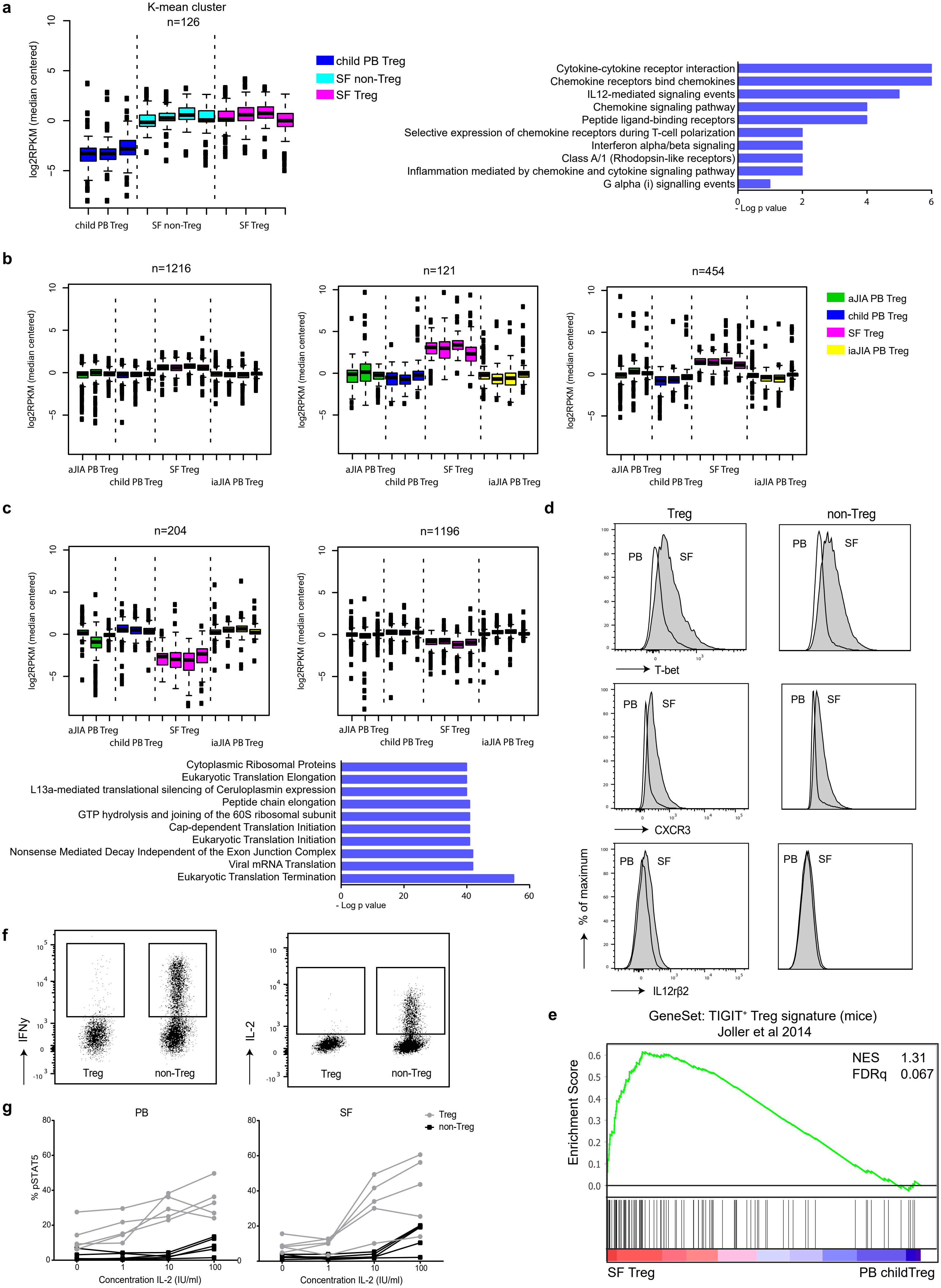

### Fig S5

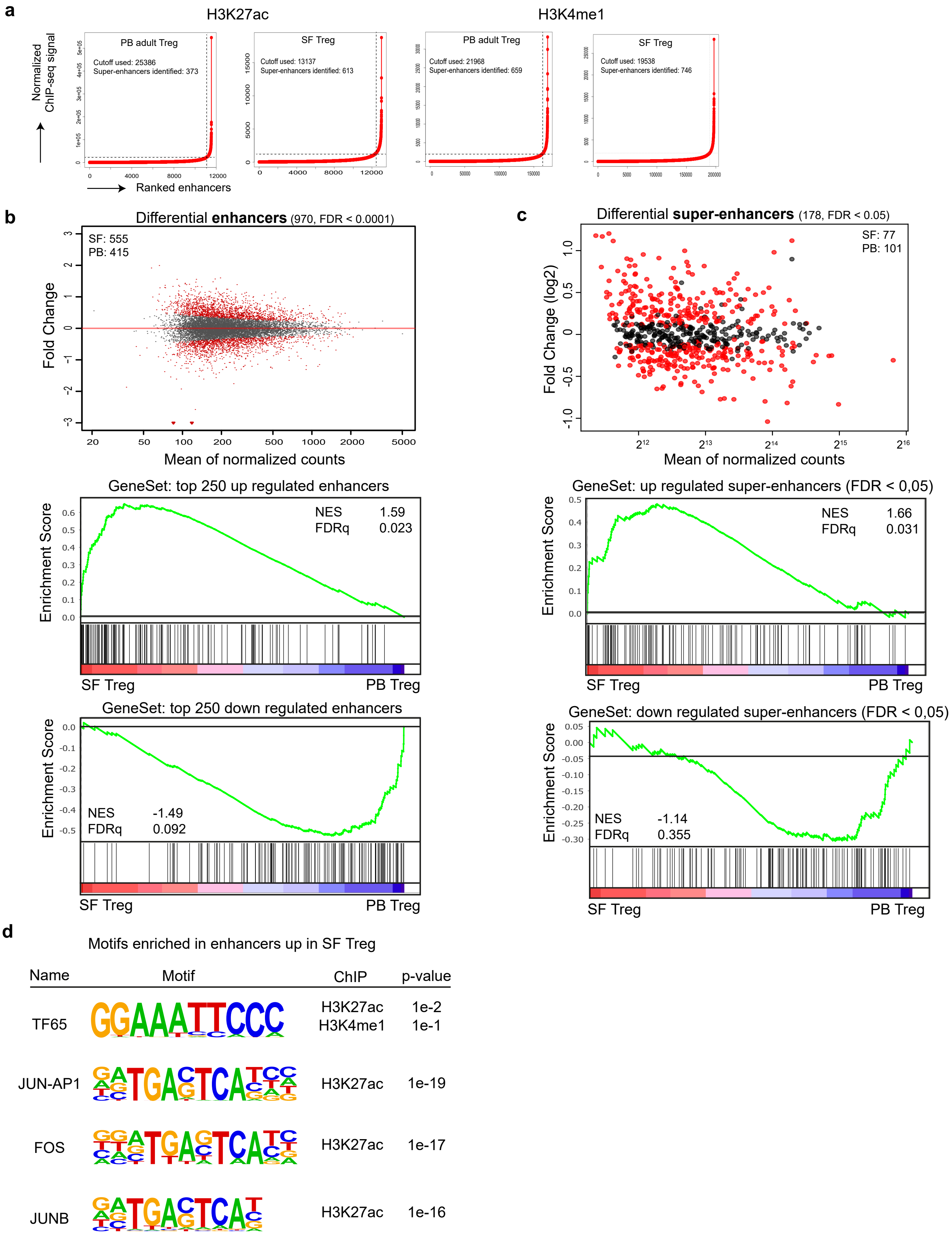

### Fig S6

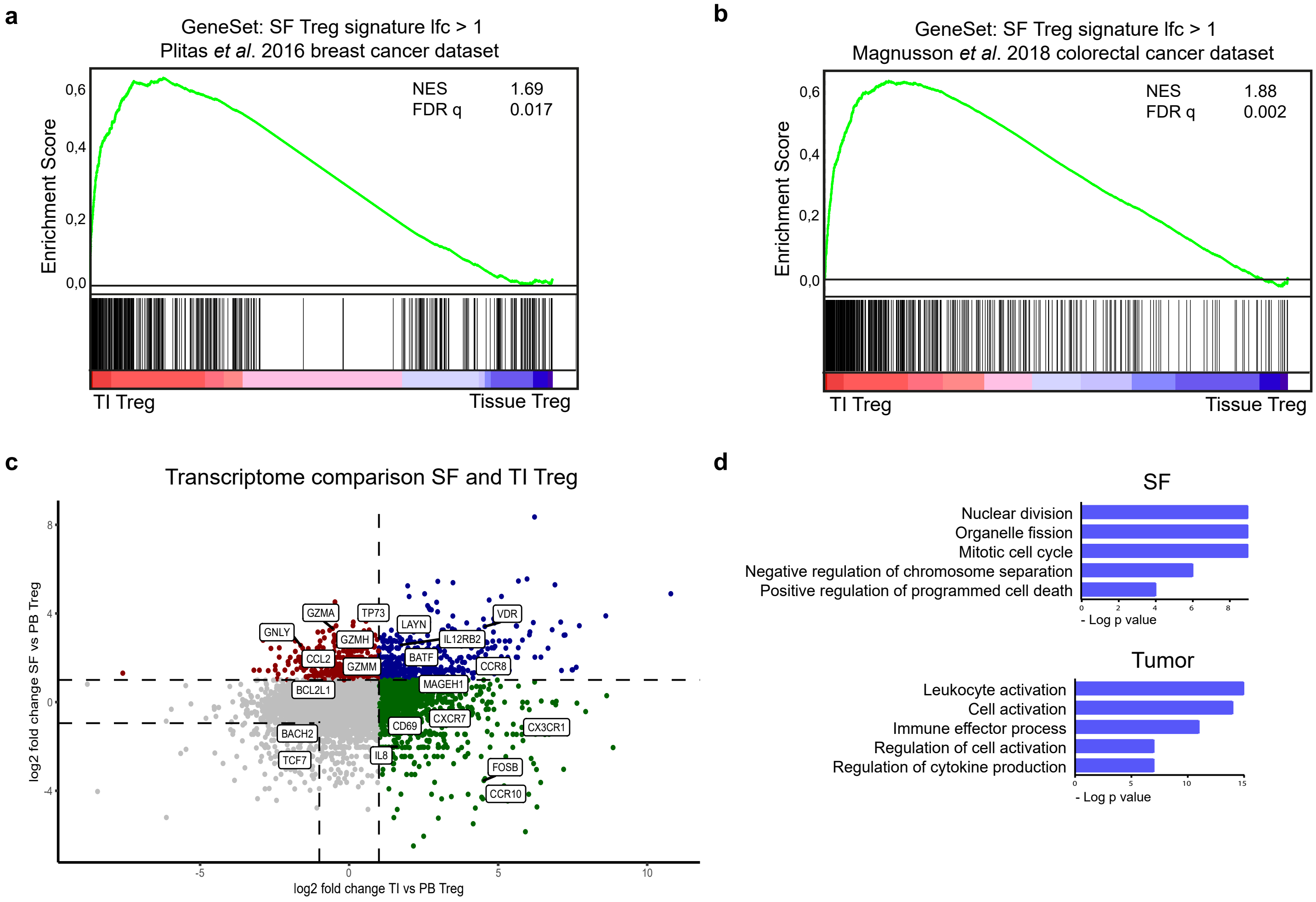
