## Supplementary material for "Conserved human effector Treg signature is reflected in transcriptomic and epigenetic landscape": Sup text

**Supplementary materials Mijnheer *et al.***

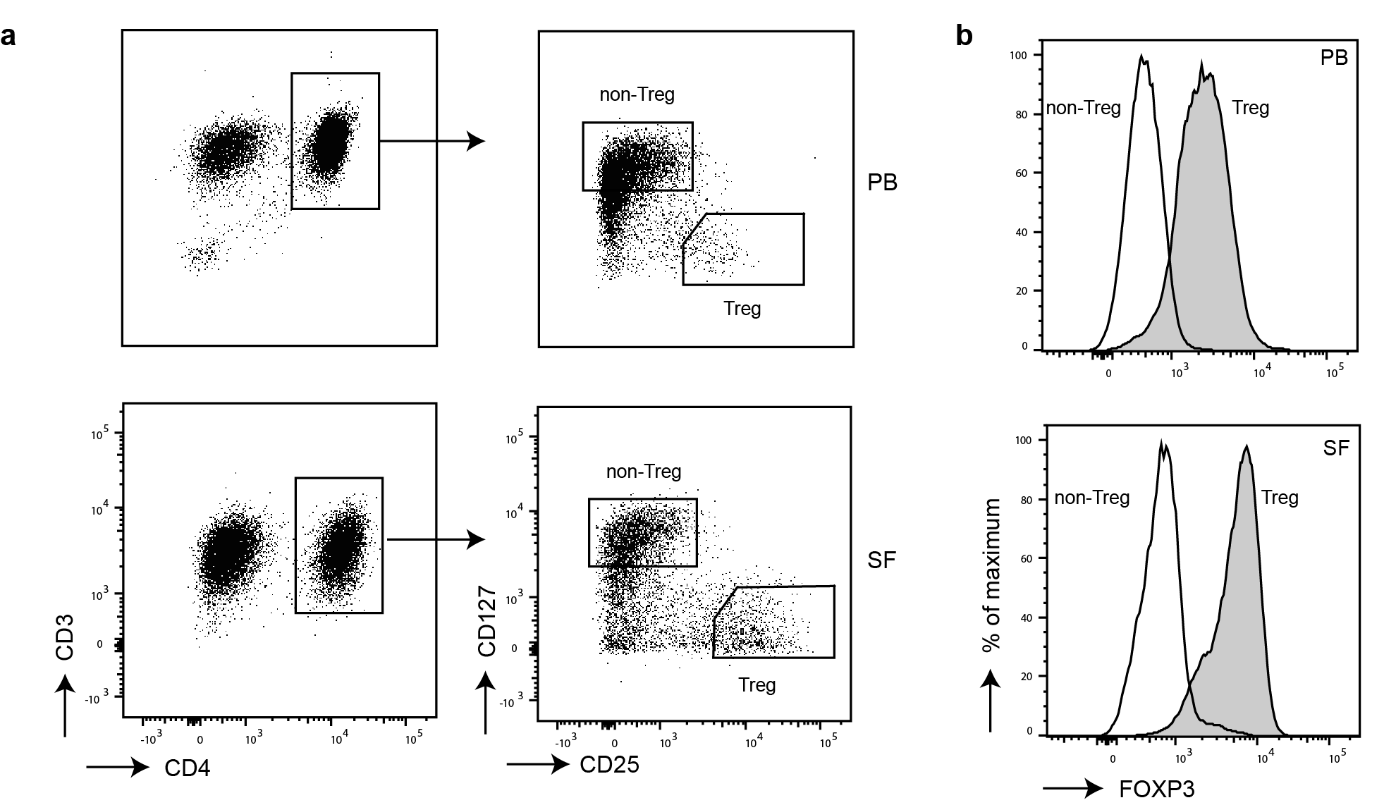

**Supplementary Figure 1.** Gating strategy and FOXP3 expression of sorted cells. **a** Representative gating strategy to sort both peripheral blood (PB)- and synovial fluid (SF)-derived Treg and non-Treg. **b** FOXP3 MFI plots of sorted SF- and PB-derived Treg and non-Treg.

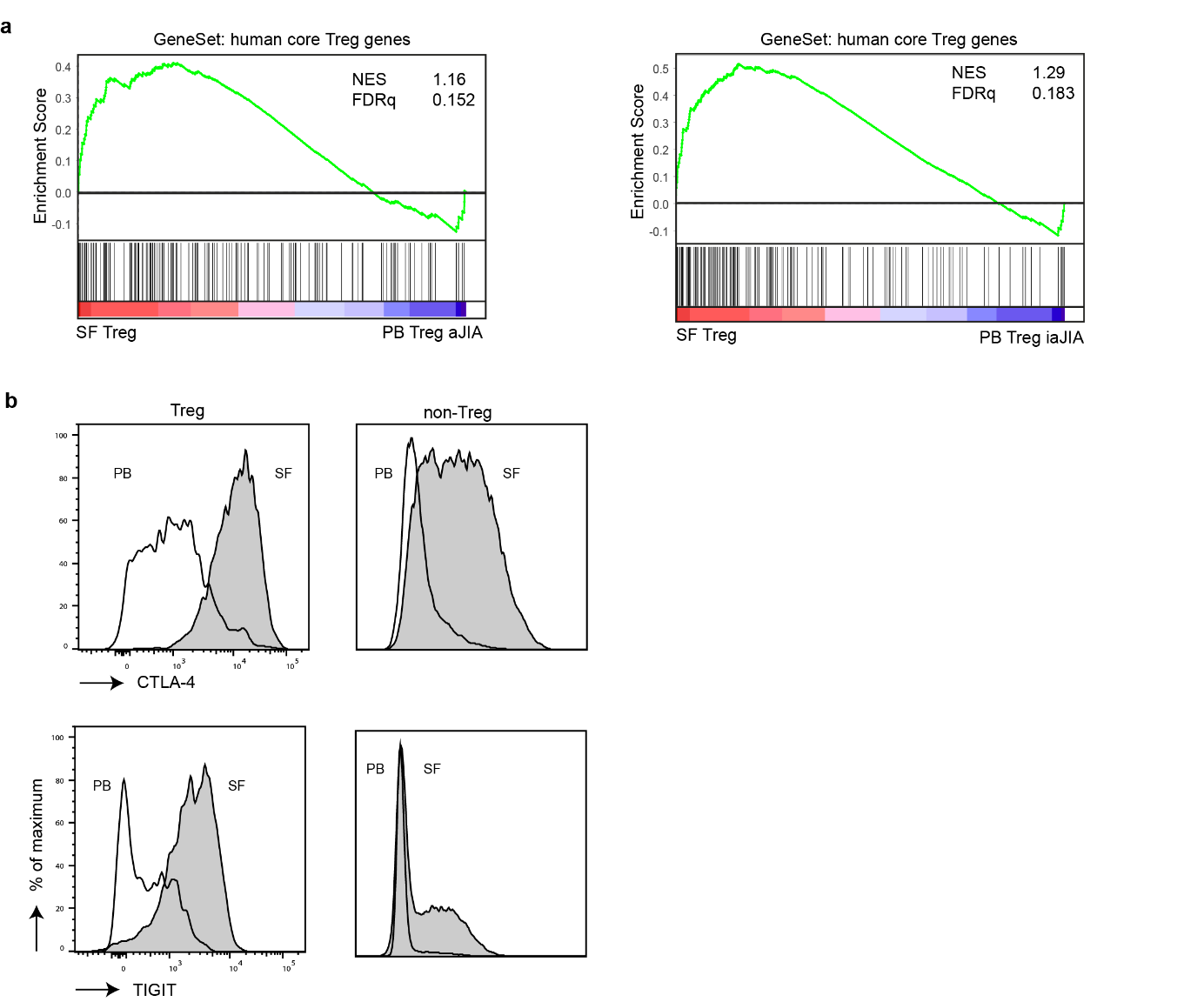

**Supplementary Figure 2.** GSEA and protein expression shows the Treg signature enriched in SF Treg. **a** GeneSet Enrichment Analysis (GSEA) of human core Treg signature genes (identified by Ferraro *et al*.^72^) in pairwise comparison of synovial fluid (SF) and peripheral blood (PB) Treg derived from juvenile idiopathic arthritis (JIA) patients with active (left) or inactive (right) disease, represented by the normalized enrichment score (NES) and FDR statistical value (FDRq). **b** Representative MFI plots (for Fig. 1e) for both CTLA4 (top) and TIGIT (bottom) in Treg (left) and non-Treg (right) derived from paired SFMC (grey fill) and PBMC of JIA patients.

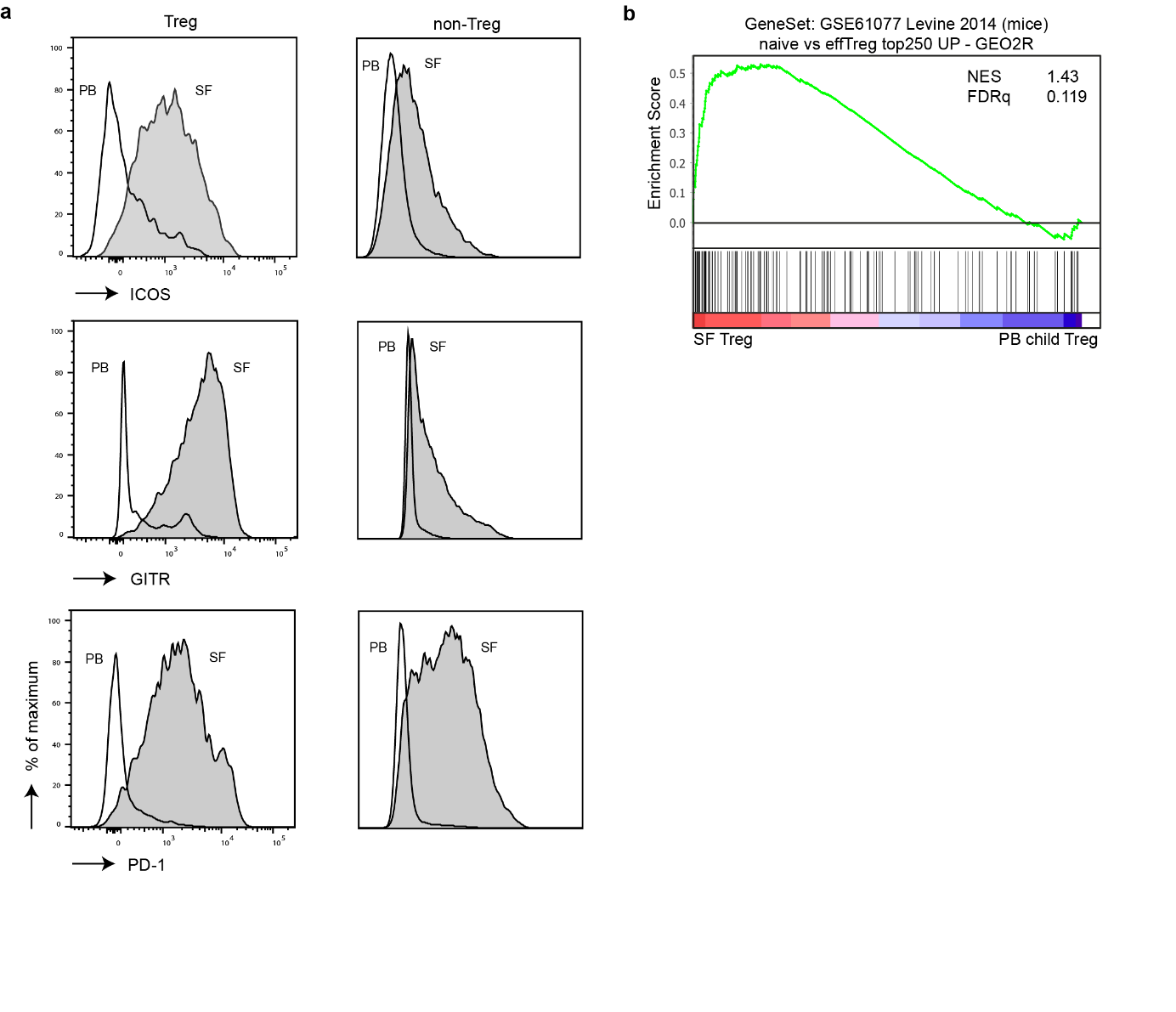

**Supplementary Figure 3.** GSEA and protein expression show an effector Treg signature in SF Treg. **a** Representative MFI plots (for Fig. 2c) of ICOS (top), GITR (middle) and PD-1 (bottom) in paired synovial fluid (SF)- and peripheral blood (PB)-derived Treg (top) and non-Treg (bottom) from juvenile idiopathic arthritis (JIA) patients. **b** GeneSet Enrichment Analysis (GSEA) of effector eTreg genes (as identified with GEO2R as the top250 genes upregulated in naïve versus eTregs, published in Levine *et al.*^21^) in pairwise comparison of SF and healthy child PB Treg, represented by the normalized enrichment score (NES) and FDR statistical value (FDRq).

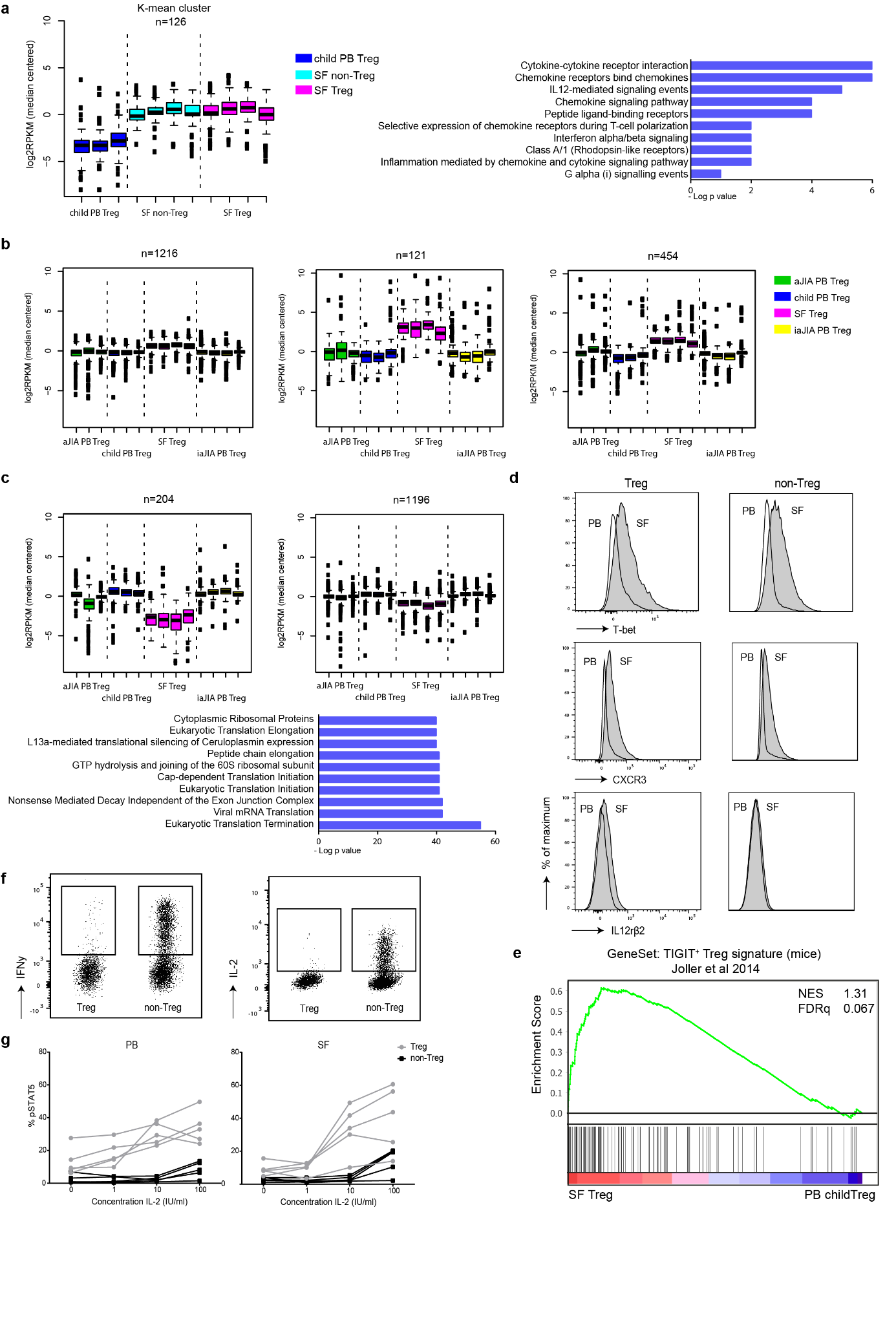

**Supplementary Figure 4.** K-mean and GeneSet Enrichment analysis shows adaptation of SF Treg to Th1 environment. **a** K-mean analysis (k = 14) on peripheral blood (PB) Treg, synovial fluid (SF) Treg and SF non-Treg with the cluster representing the genes upregulated in both SF Treg and non-Treg (n is the number of genes in the respective K-mean cluster; left) and the gene ontology pathway terms related to this cluster (right; top 10 ranked on p-value). **b** K-mean analysis (k=14) on all Treg groups derived from children (PB Treg from healthy children, juvenile idiopathic arthritis (JIA) patients with active or inactive disease and SF Treg from JIA patients) with the K-mean clusters shown representing upregulated genes in SF Treg compared to the rest(left). **c** Similar as in **b** but concerning downregulated genes in SF Treg (top) and gene ontology pathway terms (bottom; top 10 ranked on p-value). **d** Representative MFI plots (for Fig. 3d) of T-bet (top), CXCR3 (middle) and IL12rβ2 (bottom) in Treg (left) and non-Treg (right), derived from paired SFMC (grey fill) and PBMC of JIA patients. **e** GeneSet Enrichment Analysis (GSEA) of TIGIT^+^ Treg signature genes (identified in mice by Joller *et al.*^20^) in pairwise comparisons involving SF and healthy child PB Treg, represented by the normalized enrichment score (NES) and FDR statistical value (FDRq). **f** Representative dotplots (for Fig. 3f) of IFNγ (left) and IL-2 (right) in SF Treg and non-Treg. **g** Percentage of pSTAT5^+^ cells in PB- and SF-derived Treg and non-Treg in presence of increasing IL-2 concentrations (n = 5). Data are representative of two independent experiments.

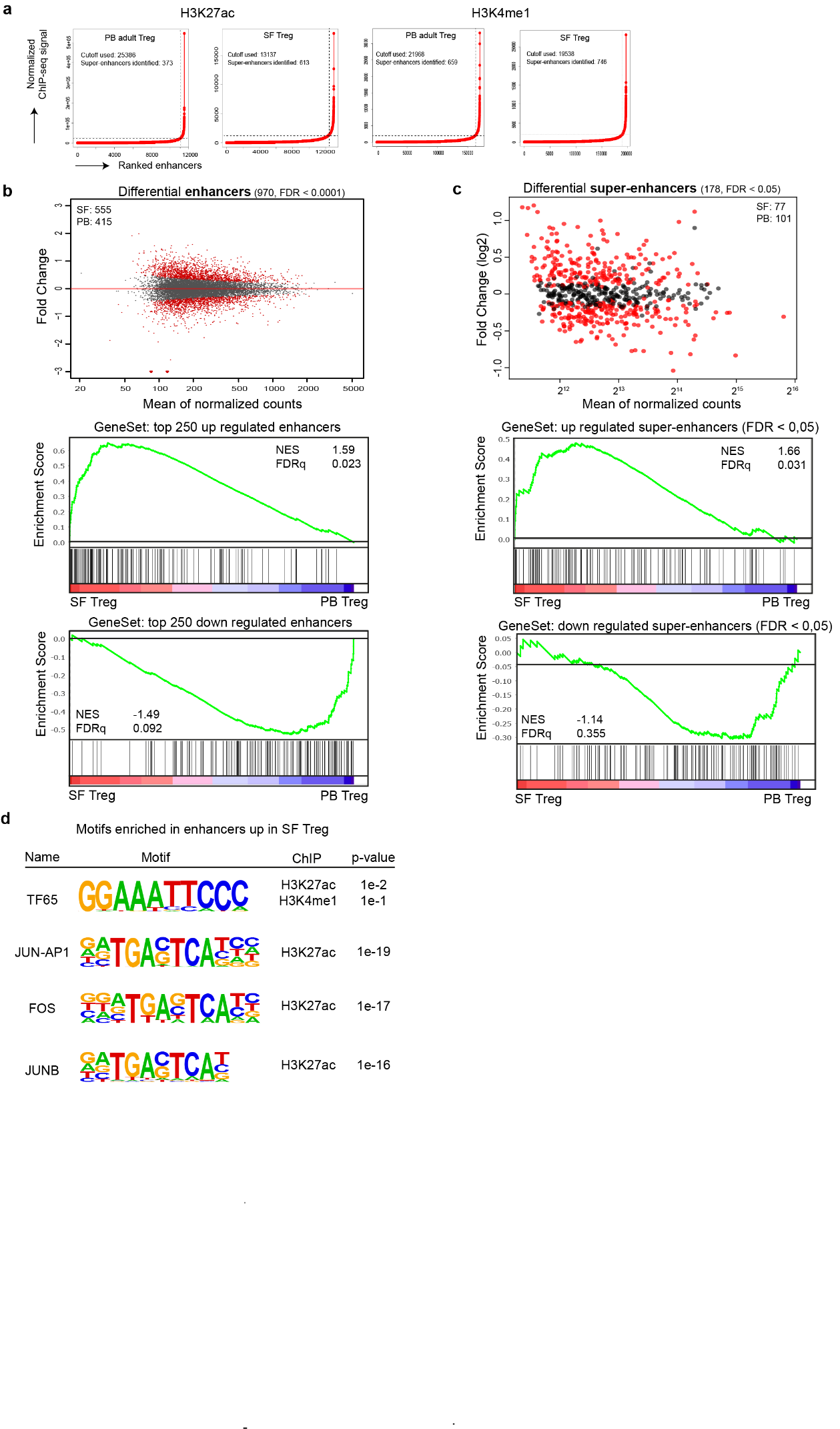

**Supplementary Figure 5.** Environment-specific effector Treg profile is regulated by the (super)enhancer landscape. **a** Representative examples of the normalized distribution of H3K27ac (left two panels) and H3K4me1 (right two panels) ChIP-seq. The plots show the enhancers ordered on the respective histone marker signal performed with the ROSE algorithm; a line with a slope of one tangent to the curve is used as a cutoff to distinguish super‐enhancers above and typical enhancers below the point of tangency. **b** MA plots of differentially expressed enhancers derived from H3K4me1 ChIP-seq (FDR < 0.0001) in synovial fluid (SF) versus peripheral blood (PB) Treg with the number of SF- and PB-specific enhancers indicated (top). GeneSet Enrichment Analysis (GSEA) of the top 250 upregulated (middle) and downregulated (bottom) enhancers in pairwise comparisons involving transcriptome data of SF Treg and PB Treg derived from healthy adults, represented by the normalized enrichment score (NES) and the FDR statistical value (FDRq). **c** Same as in **a** but for super-enhancers (FDR < 0.05). **d** Motifs, known and *de novo*, for transcription factor binding sites predicted using HOMER, enriched in the upregulated (super-)enhancers in SF Treg compared to healthy adult PB Treg for H3K27ac and H3K4me1 ChIP-seq.

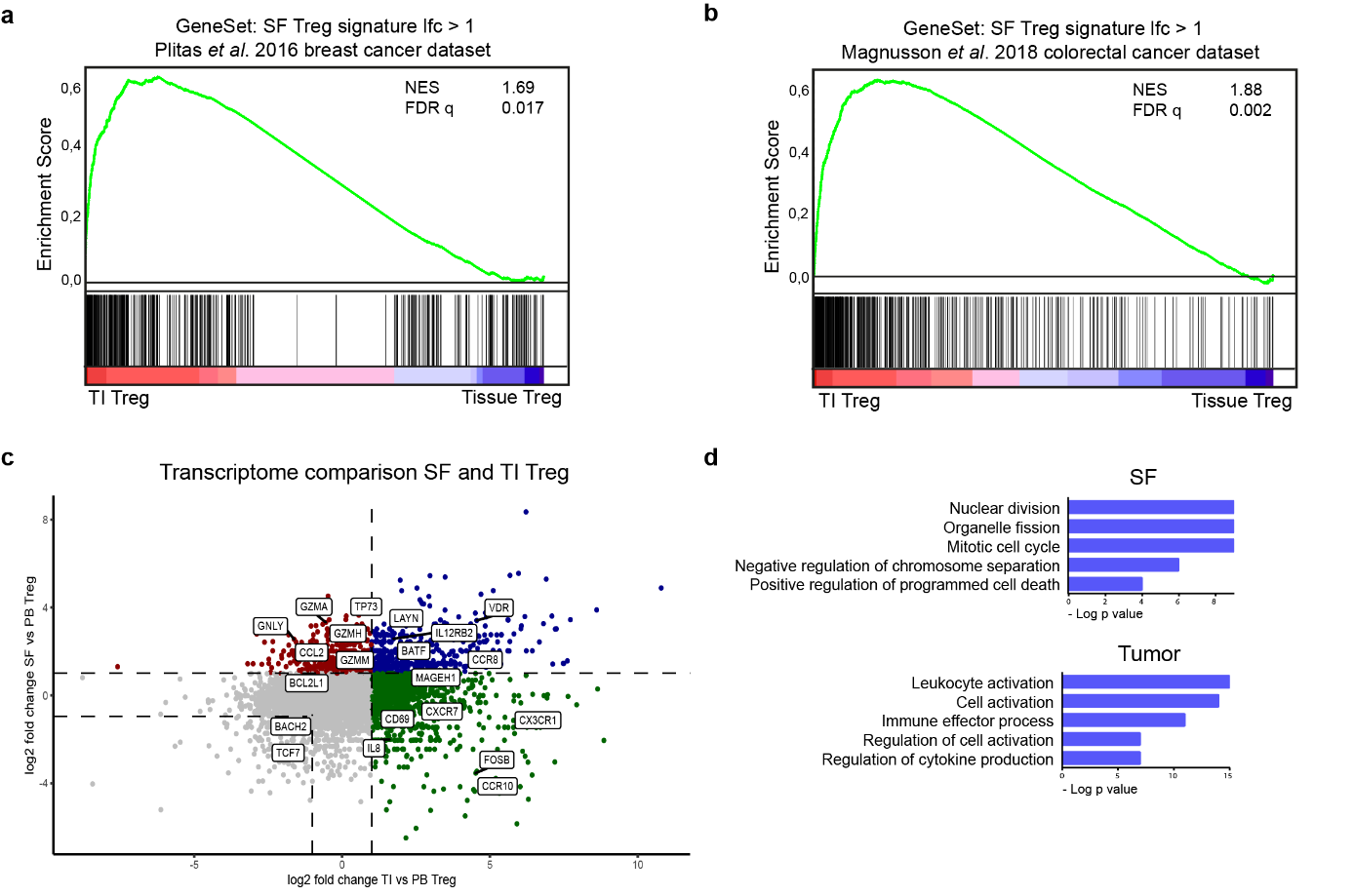

**Supplementary Figure 6.** The effector Treg program is universal and overlaps with the human tumor Treg signature. **a** GeneSet Enrichment Analysis (GSEA) of the differentially expressed genes (log2 fold change (lfc) > 1, FDR < 0.05) in synovial fluid (SF) compared to peripheral blood (PB) Treg in pairwise comparisons involving tumor-infiltrating (TI) Treg and normal tissue Treg derived from breast cancer tissue (Plitas *et al*.^16^), represented by the normalized enrichment score (NES) and FDR statistical value (FDRq). **b** Same as in **A** but concerning colorectal cancer (Magnuson *et al*.^17^). **c** Comparison of transcriptomes between SF and PB Treg to TI and normal tissue Treg. The log2 fold change of SF to PB Treg (y-axis) versus TI to PB Treg (x-axis; derived from Plitas *et al.*^16^) is shown. Each dot represents a gene present in both datasets; in grey unchanged/downregulated in both effector Treg subsets compared to PB Treg, in blue upregulated in both SF and TI Treg, in red upregulated in SF Treg and in green upregulated in TI Treg; selected genes are highlighted. **d** Gene ontology biological process terms related to genes specifically upregulated in SF (left) or TI (right) Treg from Plitas *et al.*^16^ compared to PB Treg in the respective datasets (healthy adults), ranked by enrichment scores.

Supplementary Table 1. Differentially expressed genes RNA-seq

Supplementary Table 2. Differentially expressed enhancers and super-enhancers ChIP-seq

Supplementary Table 3. Key-regulator analysis RNA-seq

Supplementary Table 4. Quality control data RNA- and ChIP-seq
